# Integrating simulated and experimental data to identify mitochondrial bioenergetic defects in Parkinson’s Disease models

**DOI:** 10.1101/2025.04.29.651221

**Authors:** Sandeep Chenna, Alvin Joselin, Daniele Bano, Paola Pizzo, Maria Ankarcrona, David Park, Jochen H. Prehn, Niamh M. C. Connolly

## Abstract

Mitochondrial bioenergetics are vital for ATP production and are associated with several diseases, including Parkinson’s Disease. Here, we simulated a computational model of mitochondrial ATP production to interrogate mitochondrial bioenergetics under physiological and pathophysiological conditions, and provide a data resource that can be used to interpret mitochondrial bioenergetics experiments. We first characterised the impact of several common respiratory chain impairments on experimentally-observable bioenergetic parameters. We then established an analysis pipeline to integrate simulations with experimental data and predict the molecular defects underlying experimental bioenergetic phenotypes. We applied the pipeline to data from Parkinson’s Disease models. We verified that the impaired bioenergetic profile previously measured in *Parkin* knockout neurons can be explained by increased mitochondrial uncoupling. We then generated primary cortical neurons from a *Pink1* KO mouse model of Parkinson’s, and measured reduced OCR capacity and increased resistance to Complex III inhibition. Here, our pipeline predicted that multiple respiratory chain impairments are required to explain this bioenergetic phenotype. Finally, we provide all simulated data as a user-friendly resource that can be used to interpret mitochondrial bioenergetics experiments, predict underlying molecular defects, and inform experimental design.

**Highlights:**

- The complexity of mitochondrial bioenergetics can make experimental data difficult to interpret.
- We simulated a computational model of mitochondrial bioenergetics in healthy and pathological conditions, and established an analysis pipeline to integrate model simulations with experimental data.
- We applied the pipeline to data from Parkinson’s Disease models to predict the molecular defects underlying Parkinson’s-related pathology.
- We provide all outputs in a user-friendly Excel file, which serves as a valuable resource to the community for insight into the effects of pathology on mitochondrial bioenergetics and for interpretation of experimental results.

## Introduction

Mitochondrial ATP production (oxidative phosphorylation) is achieved through the oxygen-dependent maintenance of a proton circuit by the mitochondrial respiratory chain (electron transport chain), the F_1_F_o_ ATP synthase, and proton leaks (Nicholls, 2013). The mitochondrial respiratory chain is a series of four multi-subunit complexes (complexes I-IV) embedded in the inner mitochondrial membrane. These complexes, via the pumping of protons (H^+^) out of the mitochondrial matrix coupled with electron transport through the complexes, serve to generate and maintain the electrochemical proton-motive force (Δ*p*), composed of the H+ concentration gradient (ΔpH_m_) and the mitochondrial membrane potential (ΔΨ_m_). This gradient drives ATP synthesis by enabling H+ flow back into the matrix through the F_1_F_o_ ATP synthase, and also maintains metabolite transport and ion homeostasis. H+ can also flow into the matrix through H+ leaks, bypassing the F_1_F_o_ ATP synthase and contributing to mitochondrial uncoupling, the dissociation between electron transport and its use to drive ATP synthesis (Brand and Nicholls, 2011, Divakaruni and Brand, 2011). Substrates generated by the tricarboxylic acid (TCA) cycle, such as NADH and FADH, serve as electron donors to the respiratory chain.

Altered mitochondrial bioenergetics have been associated with a wide variety of pathologies, including in most neurodegenerative diseases (NDs), and often emerge prior to clinical symptoms (Lin and Beal, 2006, Moran et al., 2012, Wang et al., 2020, Malpartida et al., 2021). In this regard, neurons are particularly susceptible to disruptions in mitochondrial ATP production, as synaptic activity relies heavily on OxPhos for ATP supply (Kann and Kovacs, 2007). For example, deficiency in the expression and activity of Complex I is commonly reported in Parkinson’s (Schapira et al., 1990, Haas et al., 1995, Schapira, 2008, Parker et al., 2008), following the original observation of Parkinsonian symptoms in a group of people exposed to the Complex I inhibitor MPTP (Langston et al., 1983, Nicklas et al., 1985). In Alzheimer’s, dysfunctions of the F_1_F_o_ ATP synthase and other respiratory complexes have been reported (Patro et al., 2021), as well as reduced glycolysis, substrate supply and TCA cycle capacity (Yin et al., 2014, Theurey et al., 2019, Rossi et al., 2020). Despite expanding knowledge, an understanding of the cause and effect of mitochondrial bioenergetic dysfunction in pathology is lacking.

Mitochondrial bioenergetic function can be evaluated experimentally through measurement of key parameters such as oxygen consumption rate (OCR), Δψ_m_, mitochondrial NAD(P)H or ATP (Connolly et al., 2018, Schmidt et al., 2021), and the response of these parameters to specific inhibitors, such as Rotenone or Antimycin A (to inhibit complex I and complex III respectively), provides insights into the function of distinct segments of the system. These parameters are also disrupted in cellular and animal models of NDs e.g., (Amo et al., 2011, Dong et al., 2019, Theurey et al., 2019, Park et al., 2020). Given the intrinsic complexity of mitochondrial bioenergetics, however, experiments require careful interpretation (Schmidt et al., 2021) and elucidating the molecular causes underlying bioenergetic dysfunction requires strong knowledge of mitochondrial physiology. Measurement of decreased maximal respiratory capacity in respirometry experiments, for instance, could be caused by an underlying defect in any of the respiratory chain complexes, a reduction in proton leak across the membrane (which reduces the impact of pharmacological uncoupler), or a reduced substrate supply to the respiratory chain. Optimal experimental design to pinpoint such dysfunction is vital.

Computational models, based on mechanistic details of relevant pathways, can aid in such data interpretation and experimental design, and have been used to establish causal relationships of bioenergetic defects in NDs (Cloutier and Wellstead, 2012, Bertsch et al., 2017, Chenna et al., 2021, Muddapu and Chakravarthy, 2021). Based on these considerations, we developed a computational pipeline to integrate experimental and simulated data and to predict the molecular defects underlying bioenergetic impairments in Parkinson’s Disease models. We identified reduced respiratory chain function in primary neurons from *Pink1* knockout mice, and applied the integrated pipeline to predict that a combination of mild respiratory chain defects is required to explain this bioenergetic phenotype. All simulated data are openly provided and available to aid with experimental interpretation and design.

## Methods

### Computational model of the mitochondrial respiratory chain

The computational model was adapted from a model originally developed by Beard (Beard, 2005) and further extended. The models is a flux-based, thermokinetic, ordinary differential equation model of the mitochondrial respiratory chain and ATP production (Figure 1) and is thermodynamically, mass and charge balanced. The model has been used to analyse isolated cardiac and liver mitochondria, *in vivo* skeletal muscle, intact cancer cells, and primary neurons (Wu et al., 2007, Dash et al., 2008, Huber et al., 2011, Theurey et al., 2019). The model simulating physiological conditions (PC; in the absence of any impairments) was calibrated to various measurements in non-transgenic primary cortical neurons [ref Fig1b-d from (Theurey et al., 2019)], and was recently expanded to include respiratory chain-derived reactive oxygen species (ROS) metabolism (H_2_O_2_) (Chenna et al., 2021). We have utilised a relatively simplified model to enable faster calibration, lower computational burden, and easier interpretation by experimentalists. While inclusion of additional detail may capture a broader range of biological processes, we focussed on key relationships and variables to provide accessible and meaningful explanations and predictions.

**Figure 1:**
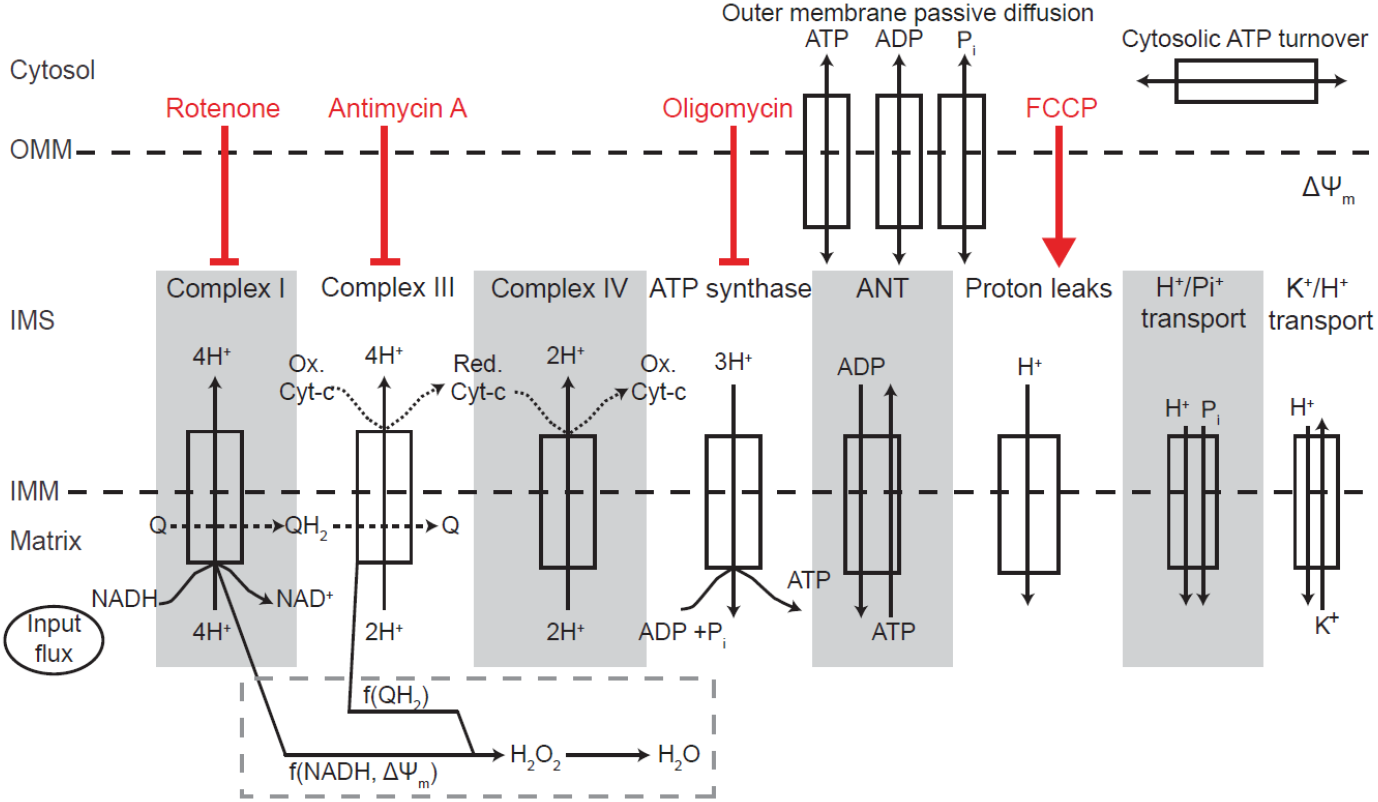
Schematic of the components and fluxes included in the computational model. Figure adapted from (Chenna et al., 2021). Addition of pharmacological modulators are simulated by inhibiting (red flat-headed arrows) or increasing (red arrow) the activity of the indicated model component, and arrows point to the text label of the affected model component. We assume the rapid conversion of superoxide to hydrogen peroxide (H_2_O_2_; not shown). ANT: adenosine nucleotide transferase; ΔΨ_m_: mitochondrial membrane potential; H^+^: protons; H_2_O_2_: Hydrogen peroxide; IMM: inner mitochondrial membrane; IMS: inter-membrane space; OMM: outer mitochondrial membrane; P_i_: Phosphate; Q: Ubiquinone; QH_2_: Ubiquinol.

The model comprises three compartments (mitochondrial matrix, inter-membrane space, cytosol) and the major model components include the respiratory chain complexes CI, CIII, CIV, the F_1_F_o_ ATP synthase (F1), proton leak across the inner mitochondrial membrane (Hle), nucleotide, ion, proton (H^+^) and substrate transport across the mitochondrial membranes, a coarse-grained model of cytosolic ATP consumption/production (Huber et al., 2011) and respiratory chain-mediated H_2_O_2_ metabolism (Figure 1). The model input is provided by a dehydrogenase flux (DH), encompassing NADH/substrate supply upstream of the mitochondrial respiratory chain. CII is considered together with CI, and CII substrates are embedded in this input dehydrogenase flux. Model outputs are represented by 22 state variables, or bioenergetic parameters, including cytosolic and mitochondrial ATP, mitochondrial membrane potential (ΔΨ_m_), mitochondrial NADH (representing redox status [NADH/NAD^+^]) and H_2_O_2_ concentrations. The rate of change of these bioenergetic parameters is defined by the flux through/activity of the modelled components. For example, the simulated NADH concentration at a given timepoint is determined by the combined activities of the NADH production reaction (model input) and the NADH consumption reaction (Complex I). The model was simulated in MATLAB (2017a).

To model the normal variability that occurs in any cell population, all simulations were performed ≥50 times with parameter values and initial concentrations varied within a normally distributed range of +/-20%, as described previously (Theurey et al., 2019). When simulating impaired complex activity (see below), sampled parameter combinations occasionally led to numerical instability (NA output). If this occurred for ≥3 simulations in a population (>5%), the parameter being adjusted was varied by the 4^th^ decimal point (e.g., 0.009 varied to 0.009001) until numerical stability was maintained.

### Data and Code Availability

The code described in this paper is freely available online at https://github.com/SandeepRed/MitoExperiments_Interpret/tree/main. Model equations, parameter values, and initial conditions are provided in Supplementary Methods. We simulated all experimental parameters described in the manuscript, in the presence of all impairments, and provide all outputs in the form of an Excel file (Supplementary Data).

### Simulating drug additions and specific experiments

Oligomycin addition (F_1_F_o_ ATP synthase inhibitor) was simulated by reduction of the x_F1_ parameter to reduce flux through the F_1_F_o_ ATP synthase (F1). FCCP (protonophore) was simulated by increasing the x_Hle_ parameter, modelling an increased proton leak across the inner mitochondrial membrane. Rotenone and Antimycin A (complex I and III inhibitors, respectively) were simulated by reducing the x_CI_ and x_CIII_ parameters. For instance, the standard ‘mitochondrial stress test’ performed in respirometry experiments to measure oxygen consumption rate (OCR) (Connolly et al., 2018) can be replicated by simulating sequential addition of Oligomycin, FCCP and Rotenone/Antimycin A. Here, oxygen consumption was assumed as the simulated flux through the oxygen-consuming CIV and the simulated OCR (mol O_2_/s/litre of mitochondria) was converted into experimental units (mol O_2_/min/μg protein) using values for mitochondrial volume (4×10^−14^ l), protein/well (45 μg), and neurons/well (300,000) as calculated previously (Theurey et al., 2019). The OCR metrics reported here are ‘basal’ (OCR in baseline conditions), oligomycin-insensitive or ‘leak’ (OCR following the simulated addition of oligomycin) and ‘maximal’ (OCR following the simulated addition of FCCP [Hle increased ∼10-fold]). Other experiments, including ΔΨ_m_, NADH, H_2_O_2_ and mitochondrial ATP changes in the presence of the aforementioned inhibitors, were simulated similarly.

### Local sensitivity analysis: Simulating impaired activity of critical model components

Sensitivity analysis measures the response of output variables to simulated changes in one or several model parameters (Saltelli et al., 2004). Sensitivity analyses on a version of this model was previously performed to rank key parameters affecting bioenergetic state variables in different conditions (*e*.*g*., healthy, apoptotic) and to map similarly behaving parameter clusters (Huber et al., 2012). To analyse the effect of specific pathology-oriented impairments on mitochondrial bioenergetic, we performed local sensitivity analysis (a single parameter is varied within a defined parameter space) and varied the activity of 8 critical model components – complex I (CI), complex III (CIII), complex IV (CIV), F_1_F_o_ ATP synthase (F1), proton leak (Hle), dehydrogenase flux (DH), cytosolic ATP production (KDyn) and cytosolic ATP consumption (KCons) – by varying their respective model parameters. Activity was varied between 2-100% of the simulated physiological condition (as calibrated to primary cortical neurons in (Theurey et al., 2019), at 2% intervals. Since component activity varies non-linearly with parameter value, we performed an exponential search to identify parameter values that yield the desired activity (Table 1).

**Table 1.** Sensitivity analysis parameter settings: Model components varied throughout this study, and the corresponding parameter values to simulate the range of impaired/increased activity. PC: simulated Physiological Condition

| Model Component (abbreviation) | Range of simulated activity (% of PC) | Model parameter | Parameter range (unitless) |
| --- | --- | --- | --- |
| Complex I (CI) | 2-100% | X<sub>CI</sub> | 0.80 - 5.75 x10<sup>-5</sup> |
| Complex III (CIII) | 2-100% | X<sub>CIII</sub> | 0.36 - 6.78 x10<sup>-4</sup> |
| Complex IV (CIV) | 2-100% | X<sub>CIV</sub> | 0.04 - 2.44 x10<sup>-5</sup> |
| F<sub>1</sub>F<sub>0</sub> ATP synthase (F1) | 2-100% | X<sub>F1</sub> | 5.50 x10<sup>-4</sup> - 3.86 x10<sup>-7</sup> |
| Proton leak (Hle) | 2-100% | X<sub>Hle</sub> | 0.98 - 1.95 x10<sup>-2</sup> |
|  | 100-1000% |  | 1 - 10 |
| Dehydrogenase flux (DH) | 2-100% | X<sub>DH</sub> | 0.75 - 7.55x10<sup>-3</sup> |
| Cytosolic ATP Production (K<sub>Dyn</sub>) | 2-100% | K<sub>ADTP_dyn</sub> | 0.96 - 9.63x10<sup>-3</sup> |
| Cytosolic ATP Consumption (K<sub>Cons</sub>) | 2-100% | K<sub>ADTP_cons</sub> | 0.98 - 1.78 x10<sup>-2</sup> |

### Categorising quantitative simulated outputs to provide a qualitative representation of model predictions

Power calculations on experimental data from healthy, non-transgenic primary cortical neurons were previously performed to define statistical thresholds beyond which predicted changes in bioenergetic variables are expected to be experimentally measurable (Theurey et al., 2019). Here, to enable similar thresholding of all model outputs, we first simulated all experiments in physiological conditions (PC) with no defect (n = 1000) and calculated the median values of all output variables. Subsequently simulated outputs (with or without a simulated impairment) were then categorised into 7 groups compared to the median PC value (Figure 2A,B). Outputs within ±10% of the median PC value were categorised as ‘no change’. Otherwise outputs were categorised as: decrease-, decrease--, decrease--- (10-25%, 25-40%, >40% decrease compared to median PC value, respectively) or increase+, increase++, increase+++ (10-25%, 25-40% >40% increase compared to median PC value, respectively) (Table 2). We estimated that outputs differing from the median by ≥25% (decrease--, decrease---, increase++, increase+++) should be detectable experimentally and that changes that do not exceed these thresholds may not be detectable experimentally. For ΔΨ_m_, NADH, H_2_O_2_ and mitochondrial ATP changes following drug additions, we also calculated the foldchange of the output over baseline (Figure 2C), as this is the commonly-reported experimental measurement (these parameters are commonly measured using fluorescent probes, and raw fluorescence measurements should not generally be compared between experiments, due to technical variability) (Scaduto and Grotyohann, 1999). For example, ΔΨ_m_ (measured in mV) is predicted to decrease in the presence of a CI impairment, and this is observed both for baseline ΔΨ_m_ and for ΔΨ_m_ following Oligomycin addition (Figure 2B). However, the ΔΨ_m_ foldchange induced by Oligomycin is not predicted to change in the presence of a CI impairment (median simulated value is ‘increase+’ compared to the simulated physiological conditions; Figure 2C).

**Table 2.**
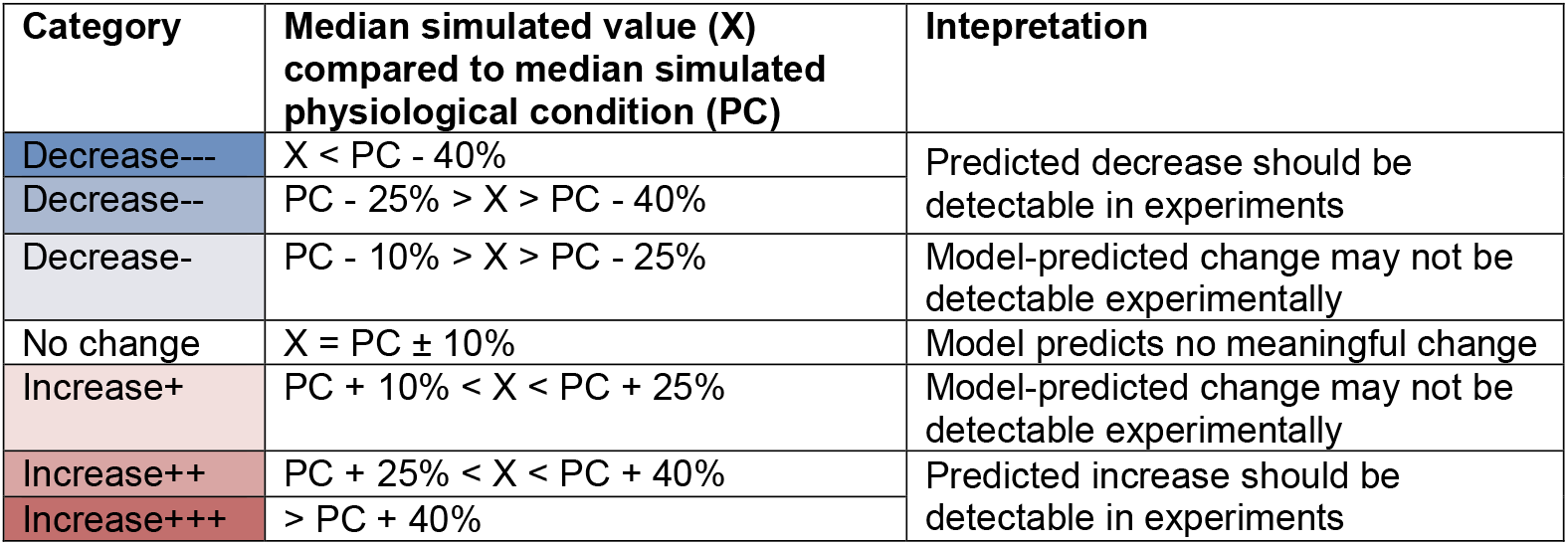
Categorising modelled outputs compared to the median of the simulated physiological condition (PC)

**Figure 2:**
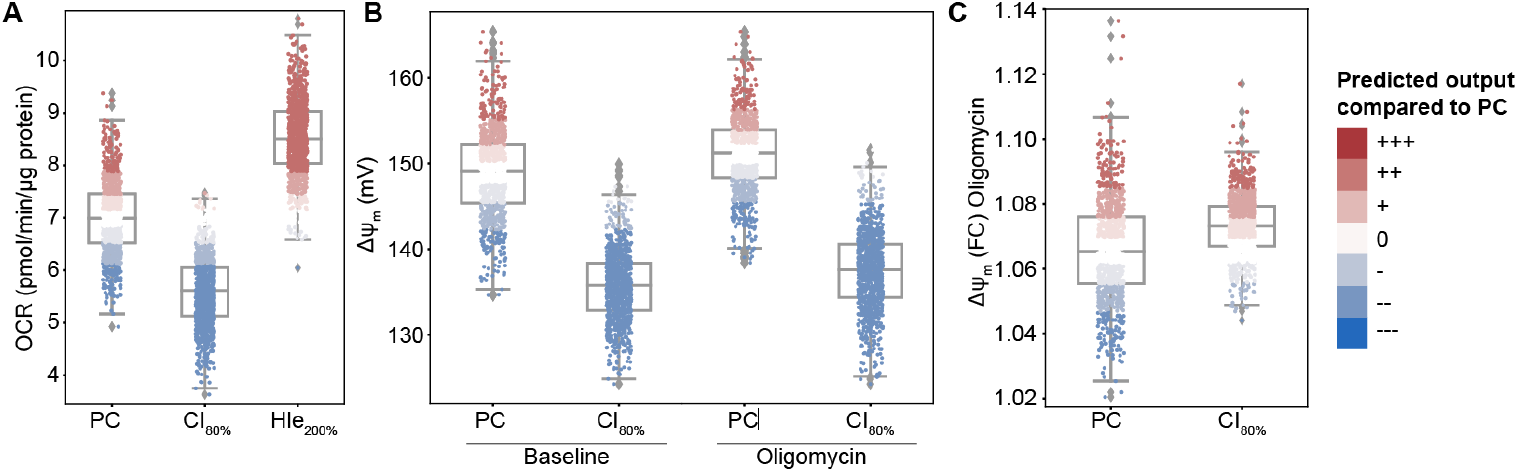
Categorising simulations provides a qualitative classification system to identify defects predicted to cause an experimentally measurable change. (A) Basal OCR simulated in physiological conditions (PC; no simulated impairment), in the presence of a complex I impairment (CI activity = 80% PC activity), and in the presence of increased proton leak (Hle activity = 200% PC activity). (B) Mitochondrial membrane potential (ΔΨ_m_) simulated at baseline and following the addition of Oligomycin (Oligo), for wildytpe conditions and in the presence of CI impairment (80% PC). (C) ΔΨ_m_ foldchange over baseline following the addition of oligomycin, in healthy and CI impairment conditions (80% PC). Outputs were grouped into 7 categories (coloured red to blue) compared to the median physiological condition: White: Median PC ± 10% (no meaningful change); Darkening shades of red indicate increase above median PC: median+10%, median+25%, median+40%. Darkening shades of blue indicate decrease below median PC: median-10%, median-25%, median-+40%. n = 1000 simulations for each condition.

### Hierarchical clustering

To identify defects that induce similar bioenergetic phenotypes, hierarchical clustering was performed in Python with the scikitlearn/seaborn libraries (Pedregosa et al., 2011, Waskom, 2021). Cluster quality was assessed by silhouette score and the separation between defects was assessed by V-measure score. The choice of linkage method did not affect cluster quality metrics in the case of qualitative inputs. For clustering of grouped data, categories were assigned numerical values (+3, +2, +1, 0, −1, −2, −3) compared to healthy simulations and clustering was performed. For clustering of both experimental and simulated data, experimental data were assigned values (+3, +2, +1, 0, −1, −2, −3) and clustering was performed identically. If the experimental phenotype clustered with the predicted phenotype of a simulated defect, this suggested that this defect may underlie the experimental model.

### Experimental methods

#### Primary cortical neurons

Experiments involving animals conformed to the guidelines set forth by the Canadian Council for the Use and Care of Animals in Research (CCAC) and the Canadian Institutes for Health Research (CIHR). All animal procedures were approved by the University of Ottawa Animal Care Committee (breeding and dissection protocol # NSI 1775 and NSI 2459, respectively). The germ line-deleted *Pink1* KO mice were obtained from Dr J. Shen and were backcrossed to C57BL/6 for more than seven generations, as described previously (Kitada et al., 2007). Primary neuronal cultures were obtained from cortices dissected from backcrossed transgenic *Pink1* (C57BL/6) KO mouse embryos at day E14.5-15.5, as previously described (Joselin et al., 2012). Experiments were performed after 8-10 days *in vitro* (DIV).

#### Seahorse respirometry

Live oxygen consumption rates (OCR) were measured using a Seahorse XF24 Analyzer (Agilent). The classical “mitochondrial stress test” protocol was performed as previously described (Connolly et al., 2018), in media including 25 mM glucose and 1 mM pyruvate. OCR was measured in 3-4 wells per condition in 3 independent preps, and normalised to protein concentration in each well, measured using a protein assay kit (Bio-Rad; Cat #500006). Non-mitochondrial OCR was subtracted. Significance was tested using a two-tailed Student’s t-test assuming equal variance.

#### TMRM fluorescence

TMRM fluorescence (10 nM) was measured in live, single neurons following standardized protocols (Connolly et al., 2018). Rotenone (2 μM) or Antimycin A (1 μM) were added after 10 minutes, and Oligomycin (2 μg/ml) was added after 30 minutes. FCCP (10 μM) was added after 70 minutes to completely depolarize the mitochondrial membrane (data not shown). Raw fluorescence intensities were normalised to baseline values (average signal intensity in the first 8 minutes prior to drug addition). Fluorescence was measured in cells from 2 independent preps, from at least n = 20 cells (10 wildtype (WT), 10 *Pink1* KO). Significance was tested after 30 minutes (prior to Oligomycin addition) using a two-tailed Student’s t-test assuming equal variance.

## Results

### Simulating impairments of critical model components provides mechanistic insight into experimentally-observed mitochondrial bioenergetic dysfunction

We aimed to develop a computational pipeline to aid in the interpretation of mitochondrial bioenergetics experiments, and to predict molecular defects underlying dysfunction. We utilised a computational model (Methods, Figure 1) that describes the mitochondrial respiratory chain (CI, CIII, CIV), mitochondrial ATP production (F_1_F_o_ ATP synthase and proton leaks), transport across mitochondrial membranes, a coarse-grained model of cytosolic ATP consumption/ production (Huber et al., 2011) and respiratory chain-mediated H_2_O_2_ metabolism (Chenna et al., 2021). We previously calibrated the model to wildtype primary cortical neurons (Theurey et al., 2019). The model input is provided by a dehydrogenase flux (DH), encompassing NADH/substrate supply upstream of the mitochondrial respiratory chain. Simulated outputs include oxygen consumption rate (OCR), cytosolic/mitochondrial ATP, mitochondrial membrane potential (ΔΨ_m_), mitochondrial NADH and H_2_O_2_ concentration.

We performed targeted local sensitivity analysis of this model to characterise the effects of respiratory chain dysfunction on key bioenergetic parameters. We simulated reduced activity of several model components, representative of processes that occur during neurodegeneration. We simulated impaired respiratory chain activity as reported in several neurodegenerative diseases (Langston et al., 1983, Yin et al., 2014, Patro et al., 2021) by reducing the activities of complex I (CI), complex III (CIII), complex IV (CIV), or the F_1_F_o_ ATP synthase (F1), and by increasing/decreasing proton leak (Hle). We also modelled impaired substrate supply by reducing the input dehydrogenase flux (DH), and defective cytosolic ATP metabolism by altering cytosolic ATP production (KDyn) or consumption (KCons). We analysed the effects of these simulated impairments on key bioenergetic parameters that are commonly measured experimentally – oxygen consumption rate (OCR), mitochondrial membrane potential (ΔΨ_m_), and basal ATP, NADH and H_2_O_2_ concentrations (Figure 3).

**Figure 3:**
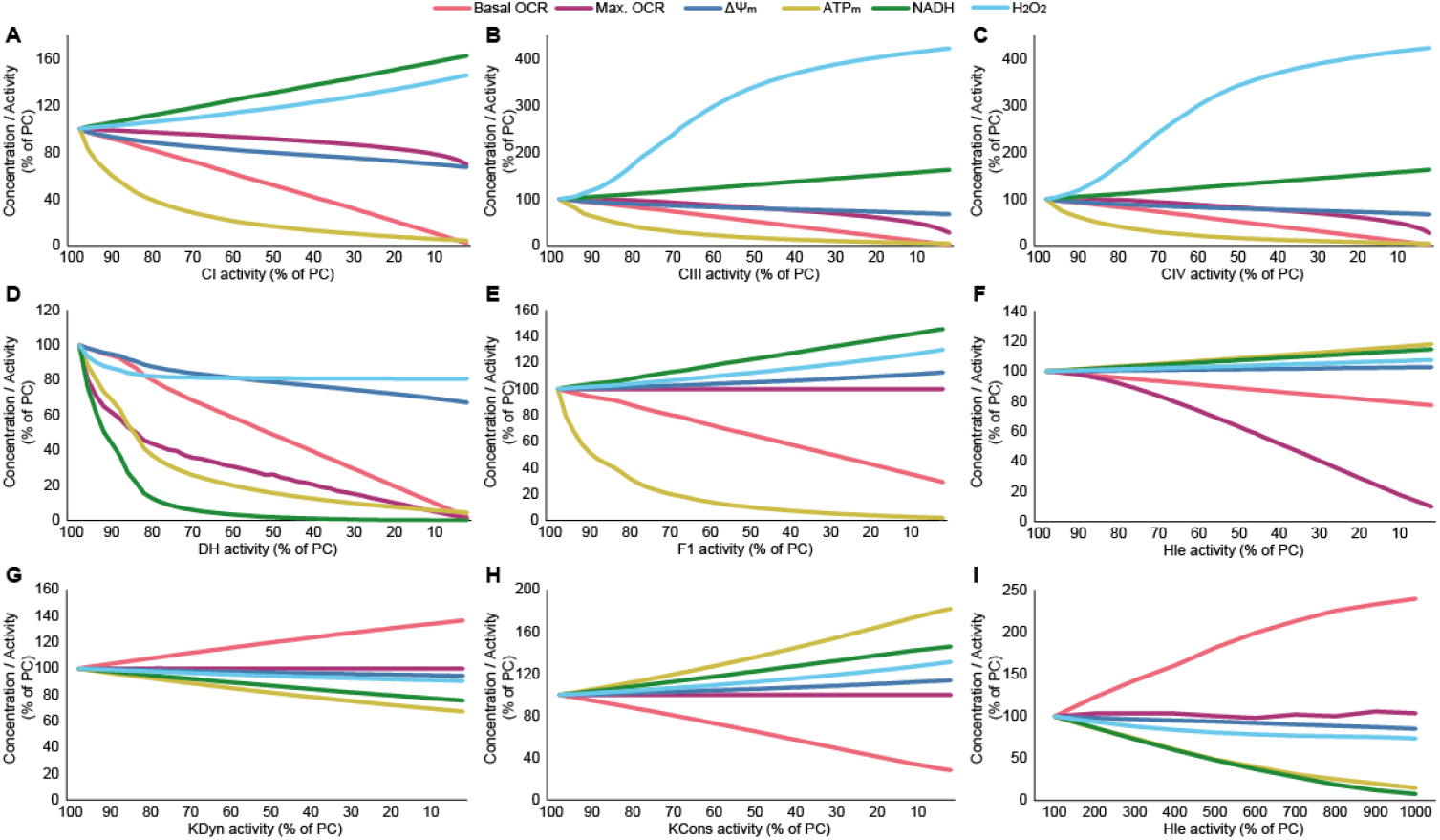
Simulating impairments of critical model components predicts their impact on bioenergetic parameters. The activity of the indicated model components (as a % of the simulated physiological conditions (PC)) is plotted on the x-axis against basal oxygen consumption rate (OCR), maximal OCR, mitochondrial membrane potential (ΔΨ_m_, normalised to a min ΔΨ_m_ of −50 mV, for visualisation), and concentrations of mitochondrial ATP, NADH, and cytosolic ROS (H_2_O_2_). Impairments were simulated in (A-C) respiratory chain complexes I, III and IV [CI, CIII, CIV]; (D) the dehydrogenase flux providing substrate to the respiratory chain [DH]; (E) the F_1_F_o_ ATP synthase [F1]; (F, I) proton leak [Hle]; (G) cytosolic ATP production [KDyn]; (H) cytosolic ATP consumption [KCons].

In the presence of these impairments, the model correctly predicted distinct behaviours based on current knowledge of cellular bioenergetics. Impairments in respiratory chain complexes (CI, CIII, CIV; Figure 3A-C) were predicted to reduce basal and maximal OCR and consequently deplete ATP_m_, while NADH levels increased due to reduced NADH consumption (Brand and Nicholls, 2011, Jafri and Kumar, 2014, Rovini et al., 2021). As respiratory complexes pump protons across the mitochondrial inner membrane to maintain the proton gradient, impaired complex activity is predicted to depolarise the mitochondrial membrane potential (ΔΨ_m_). The effect of these impairments on electron transport also increases H_2_O_2_, and the H_2_O_2_ increases predicted upon reduced CIII/CIV activity is >2-fold higher than that induced in the presence of CI impairments, as reported previously (Sipos et al., 2003, Brand, 2016, Chenna et al., 2021).

In simulations of impaired substrate supply to the respiratory chain (DH; Figure 3D), the model similarly predicted reduced basal OCR, maximal OCR and ATP_m_, and ΔΨ_m_ depolarisation. In contrast to respiratory complex inhibition, reduced DH flux substantially depletes NADH levels, as the DH flux encompasses all NADH-generating processes upstream of the respiratory chain. Simulating impaired ATP synthase activity (F1; Figure 3E) accurately predicts ATP_m_ depletion, ΔΨ_m_ hyperpolarisation [as the F_1_F_o_ ATP synthase normally consumes the proton gradient (Nicholls, 2013, Connolly et al., 2018)], and consequently a reduction in basal OCR with a corresponding NADH increase, due to decreased consumption. Maximal OCR is minimally affected by impaired F1 activity, as the capacity of the respiratory chain is driven by substrate supply and respiratory chain complex activity (Nicholls, 2013).

Proton leak (Hle) may be either increased or decreased in pathology (Divakaruni and Brand, 2011). Simulated reductions in Hle (Figure 3F) are predicted to decrease basal and maximal OCR – as maximal OCR is induced by addition of uncoupling agents such as FCCP to increase proton leakage into the mitochondria, a reduction in the simulated Hle flux alters this response. Hle reductions slightly increase NADH and ATP_m_ levels, due to increased efficiency of the respiratory chain (more of the proton gradient is available to the F_1_F_o_ ATP synthase for ATP production), and minimally affect ΔΨ_m_. In contrast, increased Hle (Figure 3I) increases basal OCR to maintain the proton gradient (although ΔΨ_m_ is still depolarised), leading to NADH depletion due to increased consumption, and ATP_m_ depletion as the proton gradient is consumed by Hle rather than available to the F_1_F_o_ ATP synthase for ATP production (Brand, 1990).

Finally, impairments in cytosolic ATP production (KDyn; Figure 3G) induce a compensatory increase in basal OCR to maintain cellular ATP, with a corresponding decrease in NADH and slight ΔΨ_m_ depolarisation. ATP_m_ is consequently depleted as more ATP is transported to the cytosol. In contrast, reductions in cytosolic ATP consumption (KCons; Figure 3H) decrease basal OCR due to reduced requirements for ATP production, with a corresponding increase in ATP_m_ and NADH concentrations, and hyperpolarisation of ΔΨ_m_. The broad effect of severe simulated impairments is summarised in Table 3, and demonstrates the accuracy of the computational model according to established knowledge of respiratory chain function. Table 3 and Figure 3 can already be used to interpret experimental data – if a decrease in NADH is measured, for instance, this may be caused by an impaired substrate supply to the respiratory chain (DH), an increase in proton leak (Hle), or impaired cytosolic ATP production (KDyn).

**Table 3.**
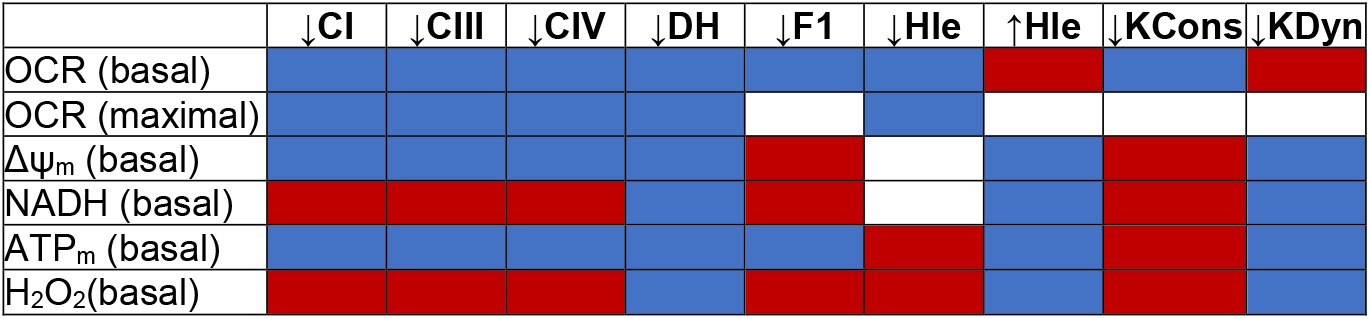
Summary of the predicted changes in bioenergetic parameters (rows) in the presence of severe impairments in the indicated model components (columns): Blue = decrease, white = no change, red = increase. Abbreviations: CI, complex I; CIII, complex III; CIV, complex IV; DH: Dehydrogenase flux, F1: F_1_F_o_ ATP Synthase, Hle: Mitochondrial proton leak, KCons: Cytosolic ATP consumption, Kdyn: Cytosolic ATP production. Down arrow indicates decreased activity of the indicated component, up arrow indicates increased activity.

### Categorising simulated responses provides a user-friendly resource to interrogate the effects of mitochondrial respiratory chain dysfunction

The addition of pharmacological agents to study mitochondrial respiratory chain function can be simulated using this model [see Methods and (Theurey et al., 2019)] and enables the replication of common experiments both in physiological conditions (PC) and in the presence of defects. Here, we visualised simulated experiments in heatmap format, enabling us to analyse, for example, the effect of Oligomycin (F_1_F_o_ ATP synthase inhibitor) on ΔΨ_m_ in physiological conditions and in the presence of CI impairments (Figure 4A). The heatmap illustrates that Oligomycin is predicted to hyperpolarise ΔΨ_m_ [foldchange (FC) > 1], but that in the presence of severe CI defects addition of Oligomycin will lead to ΔΨ_m_ depolarisation, indicating ATP synthase reversal (red box in Figure 4A). This has been demonstrated experimentally (Nicholls and Budd, 2000, Ward et al., 2000). The model also predicts, in agreement with described respiratory chain function, that Rotenone increases NADH levels due to reduced NADH consumption through CI (Rovini et al., 2021), but that this effect is smaller in the presence of a CI impairment (Figure 4B), and that Antimycin A addition similarly decreases ATP_m_ (Figure 4C; (Pauwels et al., 1985)).

**Figure 4:**
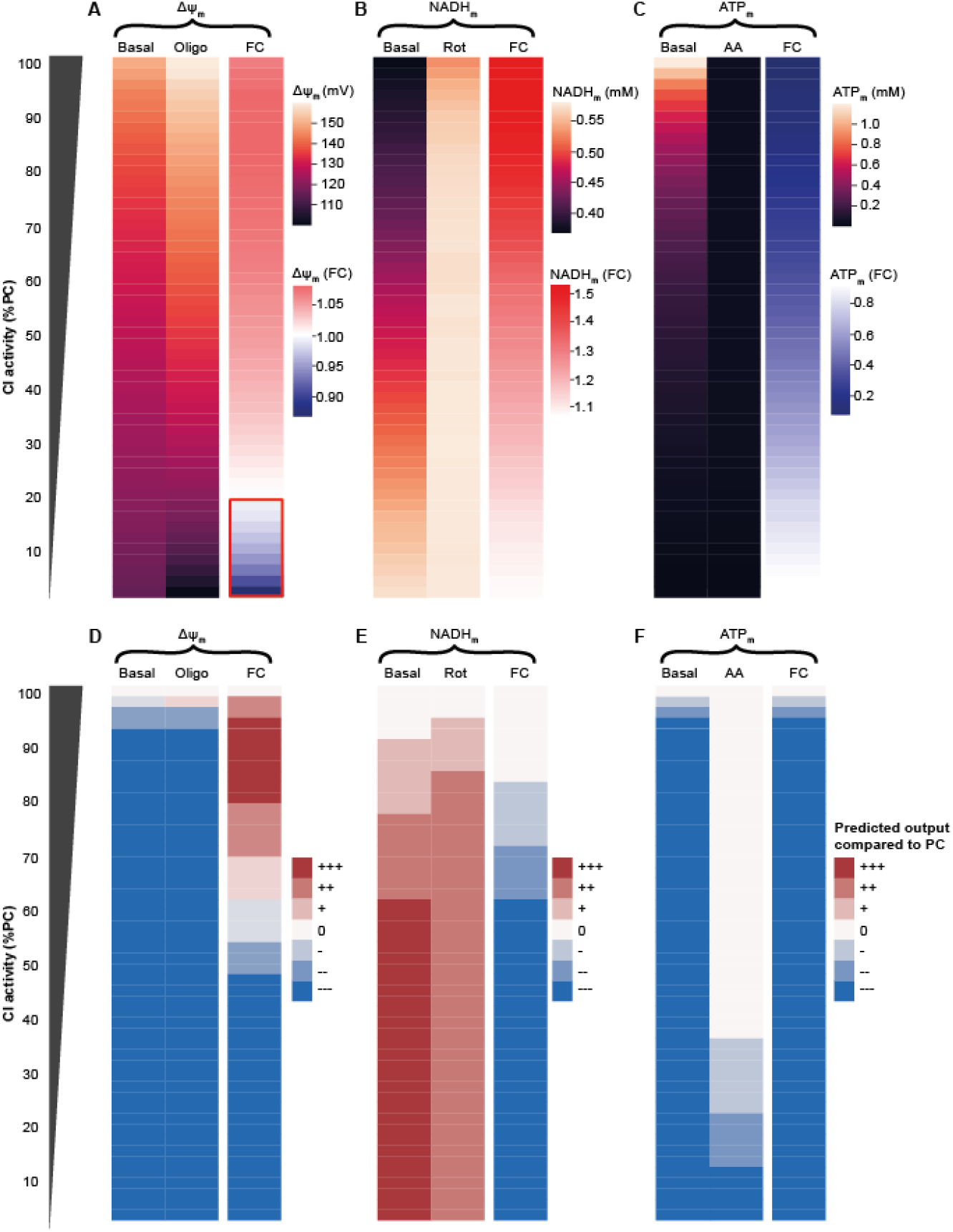
Categorising simulated impairments enables comprehensive analysis of the effects of mitochondrial respiratory chain impairments on key bioenergetic parameters. (A-C) Predicted impact of reduced complex I (CI) activity on (A) mitochondrial membrane potential (ΔΨ_m_) at baseline and following the simulated addition of Oligomycin (Oligo; F_1_F_o_ ATP synthase inhibitor), (B) mitochondrial NADH (NADH_m_) levels at baseline and following the simulated addition of Rotenone (Rot; CI inhibitor), and (C) mitochondrial ATP (ATP_m_) levels at baseline and following the simulated addition of Antimycin A (AA; CIII inhibitor). (A) The red box indicates F_1_F_o_ ATP synthase reversal predicted in the presence of severe CI defects, as evidenced by a decrease in ΔΨ_m_ following Oligomycin addition. (D-F) The same parameters are shown, with outputs grouped into 7 categories compared to the simulated physiological conditions, as described in Methods (decrease---, decrease--, decrease-, no change, increase+, increase++, increase+++). FC: foldchange (compared to basal).

To expand the utility of this resource, we grouped all simulation outputs into seven categories compared to simulated physiological conditions: decrease---, decrease--, decrease-, no change, increase+, increase++, increase+++ (see Methods). In outputs categorised as ‘no change’, ‘decrease-’, or ‘increase+’, although simulations may predict a small change in activity/concentration, we propose that this change is unlikely to be measurable experimentally. Applying these categories provides a robust qualitative demarcation and clear visualisation of the impact of defined impairments. It also aids in the interpretation of non-obvious experimental behaviour. Visualising the above-described outputs in this way (Figure 4D-F) demonstrates that in the presence of relatively mild CI impairments (≥70% of PC) the ΔΨ_m_ response to Oligo will be larger than in physiological conditions (FC increased, Figure 4D), whereas any change in the NADH_m_ response to Rotenone may not be detectable experimentally (Figure 4E). In the presence of more severe CI defects (<45% of PC) the ΔΨ_m_ response to Oligo will be smaller than that measured in physiological conditions (FC decreased, Figure 4D).

To provide a comprehensive resource to investigate the effects of mitochondrial bioenergetic impairments, we simulated all experimental parameters described in this paper, in the presence of all impairments, and provide all outputs in an Excel file (Supplementary Data).

### Unsupervised clustering enables semi-automated prediction of molecular defects underlying experimentally-observed bioenergetic phenotypes

We next combined the simulated impairments of all major model components and performed unsupervised clustering. Because impairments that induce similar bioenergetic phenotypes will cluster together, this approach enables us to visualise and identify impairments that induce similar or distinct phenotypes. For this, we analysed OCR parameters from the classical mitochondrial stress test (basal, oligomycin insensitive or ‘leak’, and maximal OCR), as well as baseline levels of ΔΨ_m_, H_2_O_2_, NADH, and mitochondrial ATP (Figure 5). We observed that impairments in F1, DH, Hle, KDyn and KCons tended to cluster separately, indicating distinct bioenergetic phenotypes induced by these impairments. Reduced DH activity, for instance, is characterised by relatively strong decreases in all parameters. In contrast, impairments in CI, CIII and CIV are predicted to induce more similar phenotypes, as flux through the respiratory chain complexes are highly correlated in unimpeded conditions (Bianchi et al., 2004), and the effects of these impairments may not be distinguishable in these settings. Nevertheless, mild defects in CI clustered away from defects in CIII and CIV, primarily due to smaller effects on H_2_O_2_ and maximal OCR. Such visualisations can be useful to quickly determine the combined effect of any impairment on the selected bioenergetic parameters.

**Figure 5:**
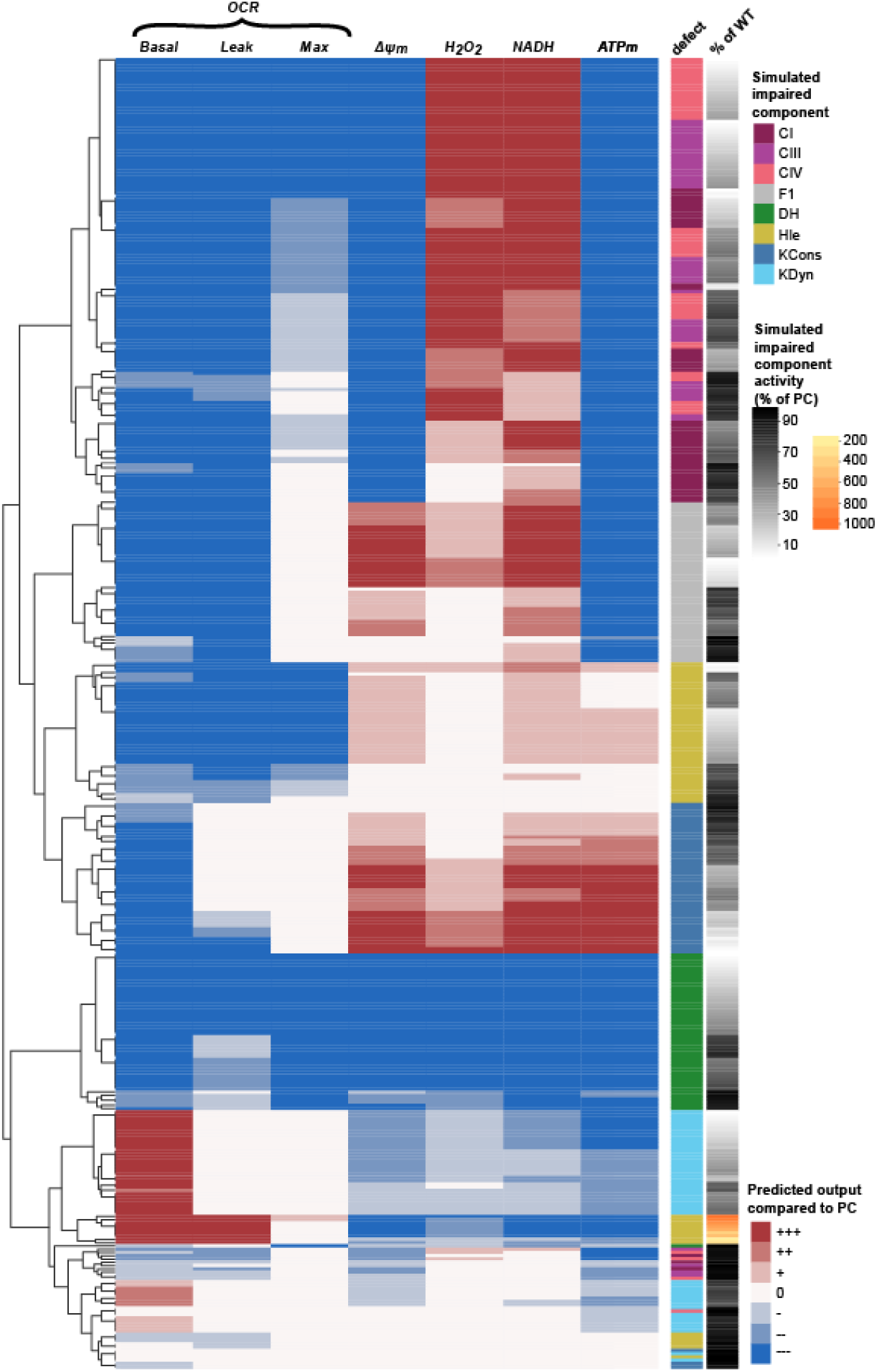
Unsupervised clustering separates impairments according to the induced bioenergetic phenotype. Impairments generally cluster together as they induce distinct bioenergetic phenotypes. Red/blue shading indicates the predicted change in the presence of the simulated impairment compared to physiological conditions (PC) as described in Methods. Row annotations indicate the component with the simulated impairment (CI, complex I; CIII, complex III; CIV, complex IV; F1, F_1_F_o_ ATP synthase; DH: dehydrogenase flux; Hle, proton leak; KCons, cytosolic ATP consumption; KDyn, cytoslic ATP production) and the magnitude of the defect [black-white: 100%-0% of activity in PC]. Leak = oligomycin-insensitive oxygen consumption rate (OCR).

### Integrated analysis verifies that a bioenergetic phenotype in Parkin knockout dopaminergic neurons can be explained by partial mitochondrial uncoupling

Disease-associated bioenergetic phenotypes are commonly reported in the scientific literature, but identifying the molecular defect underlying such phenotypes may not be straightforward. Our analysis above suggested that integrating experimentally-measured bioenergetic data (‘experimental phenotype’) into our modelling pipeline could predict the molecular defects responsible for the phenotype. By clustering the experimental data with simulated defects, we can identify which impairments produce similar bioenergetic profiles, thereby providing mechanistic insights and inferring potential underlying causes of the observed experimental phenotype. As proof of concept, we utilised experimental data in primary cortical neurons from a transgenic Alzheimer’s mouse model (Theurey et al., 2019). We integrated experimental data with simulated data in the presence of all impairments, and performed unsupervised clustering. The experimental phenotype clustered with a predicted mild defect in DH (data not shown), in agreement with subsequent experiments performed by the authors that identified reduced substrate supply to the respiratory chain (Theurey et al., 2019). This use-case confirmed our approach, that unsupervised clustering of experimental data with model simulations can pinpoint molecular defects underlying experimentally-measured bioenergetic phenotypes.

We next applied the integrated pipeline to explore the bioenergetic phenotype induced by knockout of Parkin, a Parkinson’s-related mitophagy gene known to affect mitochondrial function (Ge et al., 2020). Giguiere and colleagues previously measured increased basal OCR and unchanged maximal OCR in dopaminergic neurons from the *substantia nigra* of Parkin knockout mice, but a decrease in ATP content (Giguere et al., 2018). The authors hypothesised that this phenotype may be due to partial mitochondrial uncoupling. We clustered this experimental phenotype with simulated OCR and ATP_m_ outputs for all defects (Figure 6A). In this instance, the experimental phenotype (‘PARK’) clustered with a simulated increase in proton leak (Hle), representing increased mitochondrial uncoupling. The experimental phenotype also clustered with a simulated decrease in cytosolic ATP production (KDyn), providing an alternative molecular explanation, although severe KDyn defects (Kdyn < 20% PC) are required to reproduce the experimental phenotype. Mechanistically, these effects are explained in Figure 3I,G (increased Hle and reduced KDyn, respectively) and associated text. Indeed, subsequent simulation of OCR and ATP_m_ in the presence of increased proton leak (Hle = 350% PC) reproduced the authors’ experimental observations [compare Figure 6B with Figures 2A and 2G from (Giguere et al., 2018)], and verified that the phenotype in Parkin-knockout neurons can be explained by increased mitochondrial uncoupling.

**Figure 6:**
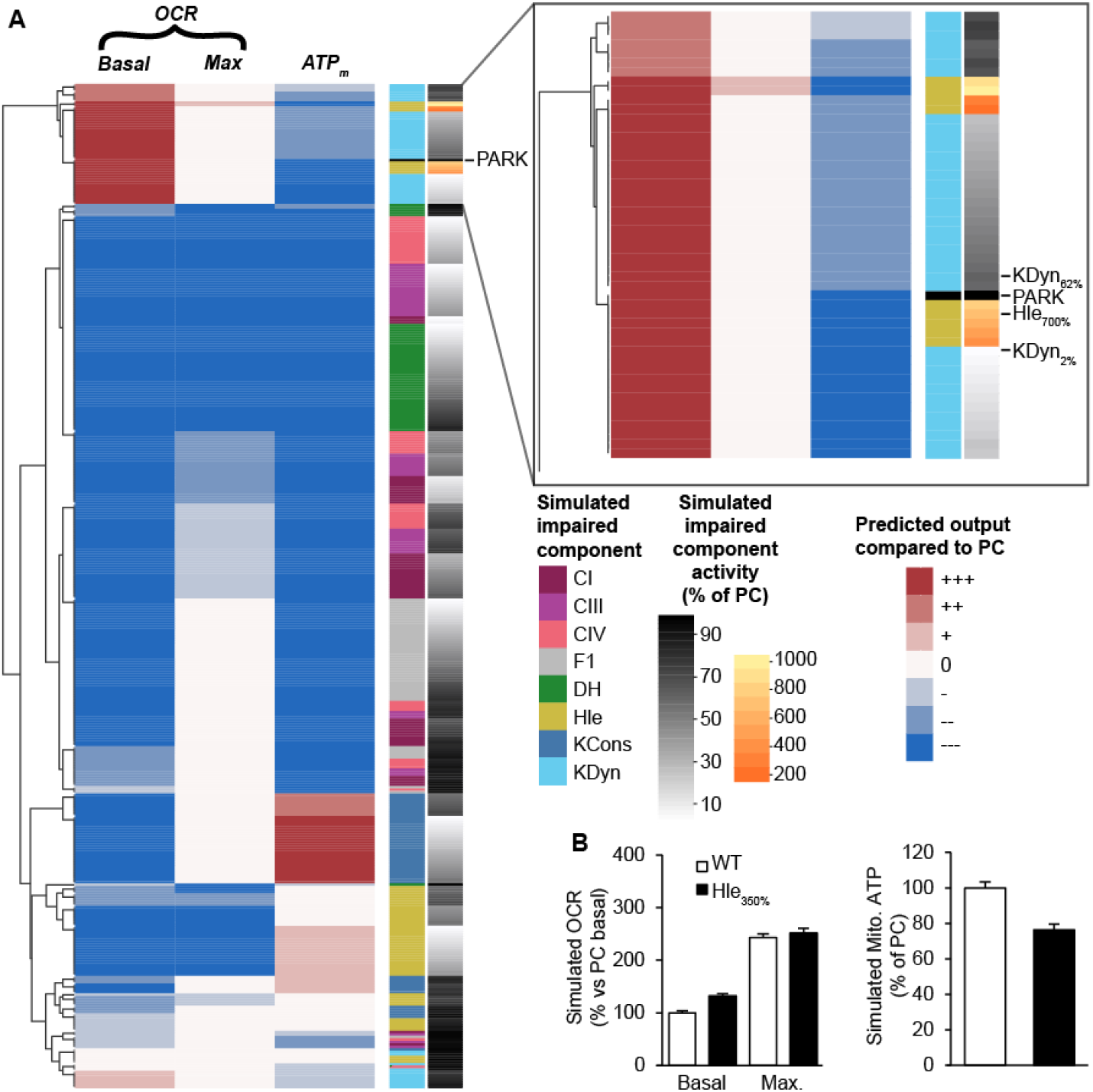
Unsupervised clustering predicts that an increased proton leak may underlie the bioenergetic phenotype measured in Parkin KO dopaminergic neurons (Giguiere et al, 2018). (A, inset) The experimental bioenergetic phenotype in Parkin KO dopaminergic neurons (PARK) clustered with a simulated increase in proton leak (Hle), equivalent to increased mitochondrial uncoupling, and large impairments in cytosolic ATP production (KDyn). Red/blue shading indicates predicted change in the presence of the simulated impairment compared to physiological conditions (PC) as described in Methods. Row annotations indicate the component with the simulated defect (CI, complex I; CIII, complex III; CIV, complex IV; F1, F_1_F_o_ ATP synthase; DH: dehydrogenase flux; Hle, proton leak; KCons, cytosolic ATP consumption; KDyn, cytosolic ATP production) and the magnitude of the defect [black-white: 100%-0% of activity in PC]. (B) A simulated increase in Hle (350% PC) reproduced the experimental observations [compare with Figures 2A and 2G from (Giguere et al., 2018)].

### De novo experiments from Pink1 knockout neurons identify impaired mitochondrial respiration, and the integrated pipeline predicts that multiple mild defects may underlie this bioenergetic phenotype

PINK1, a serine/threonine kinase localised to mitochondria, is implicated in rare inherited forms of Parkinson’s (Pilsl and Winklhofer, 2012). Along with PARKIN, PINK1 is known to play a key role in mitochondrial quality control, and knockout (KO) of the *Pink1* gene affects mitochondrial function, impairs CI activity and increases oxidative damage (Ge et al., 2020, Hoepken et al., 2007, Morais et al., 2014). We next utilised primary cortical neurons from *Pink1* KO mice to explore the effect of PINK1 on mitochondrial bioenergetic function. We measured reduced OCR across all parameters compared to WT conditions (basal, oligomycin-insensitive, maximal; Figure 7A) and a reduced sensitivity of ΔΨ_m_ to CIII inhibition (Antimycin A; Figure 7B). The response of *Pink1* KO neurons to CI inhibition (Rotenone) was similar to WT neurons (Figure 7C).

**Figure 7:**
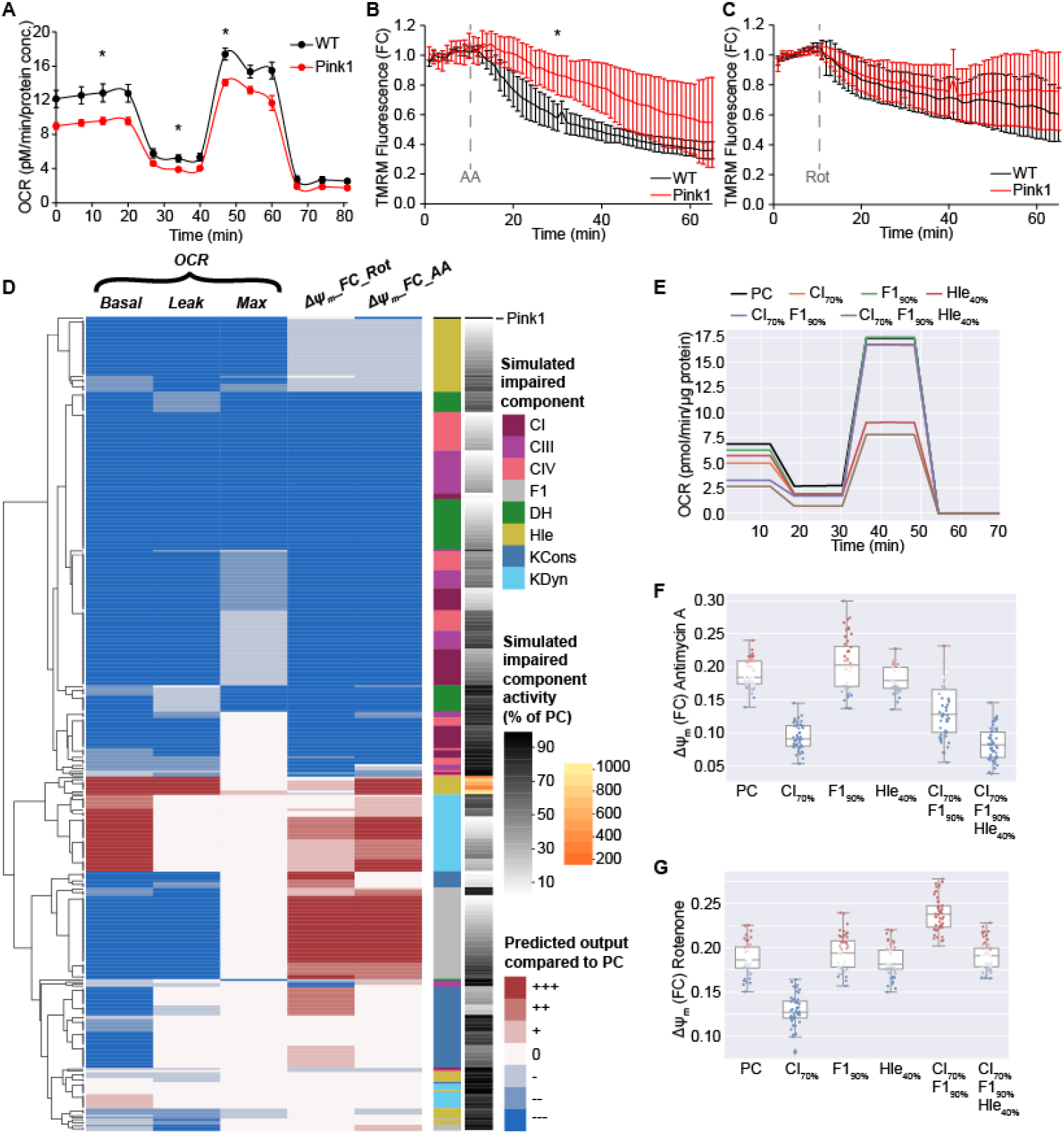
Model-guided semi-automated analysis predicts that combined defects in complex I, F_1_F_o_ ATP synthase and proton leak can mechanistically explain the bioenergetic phenotype observed in Pink1 KO neurons. (A-C) Experiments in primary cortical neurons from Pink1 KO mice identified (A) significant reductions in basal, oligomycin-sensitive and maximal oxygen consumption rates (OCR), (B) reduced ΔΨ_m_ sensitivity to Antimycin A (AA; CIII inhibition) and (C) no change in the ΔΨ_m_ response to Rotenone (Rot; CI inhibition). * p<0.05 compared to wildtype (WT). (D) Unsupervised clustering of simulated and experimental phenotypes demonstrated that while defects in Hle, CI, CIII, CIV or DH reproduced the majority of Pink1 KO (Pink1) experimental behaviour, no single defect accurately recapitulated all experiments. (E-G) Simulated experiments [(E) OCR, (F) ΔΨ_m_ foldchange (FC) response to Rotenone, (G) ΔΨ_m_ foldchange (FC) response to Antimycin A] demonstrated that the combined impairments of C1 70%PC, F1 90%PC and Hle 40%PC accurately reproduces the entire set of experiments in Pink1 KO neurons. PC; simulated physiological conditions

To explore the putative predominant molecular defect that may underlie this phenotype, we applied our integrated pipeline as above with the Pink1 KO experimental phenotype (Figure 7D). This predicted that while no single defect could explain all the observed data, several single defects explained 4/5 of the experimental observations. Specifically, a reduced proton leak (Hle <80%) is predicted to decrease all OCR metrics and does not impact the ΔΨm response to Rotenone (Figure 7D,E,G). However, in contrast to experiments, it is not predicted to impact the ΔΨm response to Antimycin A (Figure 7D,F). Similarly, defects in CI, CIII, CIV, or DH decrease OCR metrics and reproduce the decreased ΔΨ_m_ sensitivity to Antimycin A (Figure 7D-F), but these defects are also predicted to induce a strong decrease in the ΔΨ_m_ FC response to Rotenone (Figure 7D,G), an effect not observed in experiments.

We therefore hypothesised that a combination of simulated defects may be required to fully explain the experimental data. We focussed on the ΔΨ_m_ FC response to Rotenone, and reasoned that a combination of defects that both increase and decrease this response may result in the experimental observation of ‘no change’. To this end, we utilised the model to identify the defects which most strongly impact this response. We noted that defects in CI, CIII, CIV, or DH decrease the ΔΨ_m_ FC response to Rotenone, while defects in the F_1_F_o_ ATP synthase (F1) or in cytosolic ATP production (KDyn) increase the response (Figure 7D, Supplementary Data). Given that Rotenone is an inhibitor of CI, and that CI impairment can induce F1 reversal (Figure 4), we simulated the combined effect of relatively mild CI and F1 impairments, and identified that this combination significantly decreased the ΔΨ_m_ FC response to Antimycin A, while increasing the response to Rotenone (Figure 7F,G). As neither CI nor F1 defects of this magnitude reduce maximal OCR (Figure 7E), we further included a decrease in Hle, as above, and identified that defects in CI, F1 and Hle together reproduced the experimentally-observed behaviour in *Pink1* KO neurons, and may underlie this experimental phenotype (Figure 7E-G).

## Discussion

We here applied a computational model to investigate mitochondrial bioenergetic dysfunction in the presence of pathophysiological processes that occur during neurodegeneration, and established an analysis pipeline to predict underlying molecular defects that can explain experimentally-observed mitochondrial bioenergetic dysfunction. We also provide an open resource to aid interpretation of complex experimental data, provide mechanistic insight, generate hypotheses, and inform experimental design.

We first simulated several pathological impairments and analysed the predicted effect on key mitochondrial bioenergetic parameters. Simulations agreed with commonly-observed experimental data, including reduced OCR and increased NADH upon respiratory complex inhibition, mitochondrial membrane hyperpolarisation following impairment of the F_1_F_o_ ATP synthase (Jafri and Kumar, 2014, Rovini et al., 2021), and reversal of the ATP synthase in the presence of severe respiratory complex impairments (Nicholls, 2013, Connolly et al., 2018). As the model is a theoretical representation of a complex biological system, the model cannot simulate the expected biological variability as fluxes/concentrations tend to zero. We therefore simulated defects down to 2% of physiological conditions, but not beyond. These simulations demonstrated the accuracy of the model according to established knowledge of mitochondrial ATP bioenergetics, and provides precise mechanistic explanations of experimental behaviour in the presence of several bioenergetic impairments as reported in pathology. These simulations (Figure 3, Table 3) can be used as an aid to interpret mitochondrial bioenergetic experimental data.

Several computational models of respiratory chain function have been developed [*e*.*g*., (Magnus and Keizer, 1997, Cortassa et al., 2003, Berndt et al., 2015, Bazil et al., 2016, Sadri et al., 2023)]. The computational model used here simulates the respiratory chain complexes as separate entities, rather than the lumped functions in Keizer and Cortassa models (Magnus and Keizer, 1997, Cortassa et al., 2003). However, it maintains reduced complexity compared to other models [*e*.*g*., (Bazil et al., 2016, Sadri et al., 2023)] and does not incorporate, for example, beta oxidation, the effects of alternative substrates, or individual TCA cycle components. Similarly, the phenomenological ROS metabolism equations described here (Chenna et al., 2021) do not explicitly model NAD(P)H-dependent scavenging processes, which may influence redox balance and oxidative stress (Korge et al., 2015). While the inclusion of additional details may capture a broader range of biological processes, we have utilised a simplified model to enable faster calibration, lower computational burden, and easier interpretation. By focusing on key relationships and variables, our model provides meaningful and accessible explanations and predictions.

We note that the model cannot represent behaviours outside of the modelled components. A simulated impairment in the dehydrogenase flux input function, for instance, does not distinguish between TCA cycle dysfunction or defects in substrate import to the mitochondria, but rather represents a cumulative impaired supply of substrate to the respiratory chain. We acknowledge that mitochondrial bioenergetic phenotypes occur that cannot be explained by the described model system. Simulation results, as with experimental data (Schmidt et al., 2021), should be interpreted carefully.

We demonstrated that clustering simulated outputs can identify impairments that induce similar bioenergetic phenotypes. Thus, by comparing an experimental phenotype with model simulations of known respiratory chain defects, the pipeline predicts specific defect(s) that could explain the experimentally-observed phenotype. This approach enabled us to predict molecular defect(s) contributing to the bioenergetic phenotypes observed in *Parkin* and *Pink1* KO neurons. Utilising experimental data from *substantia nigra* dopaminergic neurons in a *Parkin* KO mouse model of PD (Giguere et al., 2018), the pipeline verified that the observed bioenergetic phenotype could be explained by an increase in proton leak (mitochondrial uncoupling), as had been hypothesised by the authors (Giguere et al., 2018), or by a severe decrease in cytosolic ATP production. The model also provides a mechanistic explanation for these effects (Figure 3I,G) – (i) an increased proton leak induces a compensatory increase in basal OCR to maintain the proton gradient, but this is not sufficient to maintain mitochondrial ATP levels, or (ii) reduced cytosolic ATP production leads to an increase in basal OCR to restore cytosolic ATP, but similarly is not sufficient to maintain mitochondrial ATP. To identify the more probable defect underlying the *Parkin* KO disease state, we reasoned that a severe decrease in cytosolic ATP production is less likely to occur in a biological setting. The pipeline does not predict whether this is a direct or indirect effect of *Parkin* KO, e.g. on mitophagy. A recent report suggests that any impact on respiratory chain function may be indirect (Filograna et al., 2024).

Our pipeline enables interpretation of the bioenergetic phenotype observed in a particular system. As demonstrated, this approach aims to predict the defect that induces the phenotype most similar to the experimental phenotype, rather than computing the exact solution as in a traditional linear or nonlinear least squares data fitting. This enables the generalisation of the model predictions to other experimental systems with comparable respiratory chain function (*i*.*e*. intact cultured cells respiring predominantly on CI substrates), as the direction rather than the magnitude of change is most important, and is less impacted by noise, system variability or model kinetics/parameterisation. Our pipeline also enables the ranking of putative defects, for instance by reverting to the quantitative simulated data measuring the predicted effect size of each defect.

Finally, we generated primary cortical neurons from a *Pink1* KO mouse model of Parkinson’s, and identified reduced OCR capacity and increased resistance to CIII inhibition. We applied our computational pipeline to these *de novo* data and predicted that multiple impairments may be required to completely recapitulate the experimentally-observed bioenergetic phenotype. PINK1 deficiency has been shown to impact mitochondrial bioenergetics independent of mitophagy, including decreased CI activity (Morais et al., 2014, Gautier et al., 2008), and alterations in mitophagy may also have downstream effects on bioenergetics. Disruptions in mitochondrial fission, for instance, could impair the assembly of respiratory chain complexes and lead to defective respiration (Liu et al., 2011). The predicted impairments in CI and F1 are relatively mild and may be difficult to detect in stand-alone experiments.

In conclusion, we here developed and applied a computational pipeline to interrogate mitochondrial bioenergetics in the presence of PD familial mutations. We verified that increased mitochondrial uncoupling may underlie the bioenergetic phenotype in *Parkin* KO dopaminergic neurons, and that a combination of mild respiratory chain defects is required to explain the bioenergetic impairments measured in primary cortical neurons from *Pink1* KO mice. We also provide a comprehensive resource (Supplementary Data) detailing the effects of mitochondrial respiratory chain dysfunction on key bioenergetic parameters. This resource can aid the interpretation of complex mitochondrial bioenergetic experiments, enable hypothesis generation, and inform experimental design.

## Author Contributions

**SC:** Methodology, Software, Formal Analysis, Data Curation, Writing – Original Draft, Review & Editing, Visualisation. **AJ:** Investigation, Formal Analysis. **DB:** Conceptualisation. **PP:** Conceptualisation. **MA:** Conceptualisation. **DP:** Conceptualisation, Supervision, Funding Acquisition. **JHM:** Conceptualisation, Writing – Review & Editing, Funding Acquisition. **NMCC:** Conceptualisation, Methodology, Formal Analysis, Writing – Original Draft, Review & Editing, Visualisation, Supervision.

## Ethics statement

Experiments involving animals conformed to the guidelines set forth by the Canadian Council for the Use and Care of Animals in Research (CCAC) and the Canadian Institutes for Health Research (CIHR). All animal procedures were approved by the University of Ottawa Animal Care Committee (breeding and dissection protocol # NSI 1775 and NSI 2459, respectively).

## Legends

Supplementary Data: We simulated all experimental parameters described in the manuscript, in the presence of all impairments, and provide all outputs in this Excel file.

Supplementary Methods: This Word file contains all model equations, parameter values, and initial conditions.

## References

Amo, T., Sato, S., Saiki, S., Wolf, A. M., Toyomizu, M., Gautier, C. A., Shen, J., Ohta, S. & Hattori, N. 2011. Mitochondrial membrane potential decrease caused by loss of PINK1 is not due to proton leak, but to respiratory chain defects. Neurobiol Dis, 41, 111–8.

Bazil, J. N., Beard, D. A. & Vinnakota, K. C. 2016. Catalytic coupling of oxidative phosphorylation, ATP demand, and reactive oxygen species generation. Biophysical journal, 110, 962–971.

Beard, D. A. 2005. A biophysical model of the mitochondrial respiratory system and oxidative phosphorylation. PLoS Comput Biol, 1, e36.

Berndt, N., Kann, O. & Holzhutter, H. G. 2015. Physiology-based kinetic modeling of neuronal energy metabolism unravels the molecular basis of NAD(P)H fluorescence transients. J Cereb Blood Flow Metab, 35, 1494–506.

Bertsch, M., Franchi, B., Marcello, N., Tesi, M. C. & Tosin, A. 2017. Alzheimer’s disease: a mathematical model for onset and progression. Math Med Biol, 34, 193–214.

Bianchi, C., Genova, M. L., Castelli, G. P. & Lenaz, G. 2004. The mitochondrial respiratory chain is partially organized in a supercomplex assembly: kinetic evidence using flux control analysis. Journal of Biological Chemistry, 279, 36562–36569.

Brand, M. D. 1990. The proton leak across the mitochondrial inner membrane. Biochim Biophys Acta, 1018, 128–33.

Brand, M. D. 2016. Mitochondrial generation of superoxide and hydrogen peroxide as the source of mitochondrial redox signaling. Free Radical Biology and Medicine, 100, 14–31.

Brand, M. D. & Nicholls, D. G. 2011. Assessing mitochondrial dysfunction in cells. Biochem J, 435, 297–312.

Chenna, S., Prehn, J. H. & Connolly, N. M. Phenomenological equations for electron transport chain-mediated reactive oxygen species metabolism. 2021 IEEE International Conference on Bioinformatics and Biomedicine (BIBM), 2021. IEEE, 653–658.

Cloutier, M. & Wellstead, P. 2012. Dynamic modelling of protein and oxidative metabolisms simulates the pathogenesis of Parkinson’s disease. IET Syst Biol, 6, 65–72.

Connolly, N. M. C., Theurey, P., Adam-Vizi, V., Bazan, N. G., Bernardi, P., Bolanos, J. P., Culmsee, C., Dawson, V. L., Deshmukh, M., Duchen, M. R., Dussmann, H., Fiskum, G., Galindo, M. F., Hardingham, G. E., Hardwick, J. M., Jekabsons, M. B., Jonas, E. A., Jordan, J., Lipton, S. A., Manfredi, G., Mattson, M. P., Mclaughlin, B., Methner, A., Murphy, A. N., Murphy, M. P., Nicholls, D. G., Polster, B. M., Pozzan, T., Rizzuto, R., Satrustegui, J., Slack, R. S., Swanson, R. A., Swerdlow, R. H., Will, Y., Ying, Z., Joselin, A., Gioran, A., Moreira Pinho, C., Watters, O., Salvucci, M., Llorente-Folch, I., Park, D. S., Bano, D., Ankarcrona, M., Pizzo, P. & Prehn, J. H. M. 2018. Guidelines on experimental methods to assess mitochondrial dysfunction in cellular models of neurodegenerative diseases. Cell Death Differ, 25, 542–572.

Cortassa, S., Aon, M. A., Marbán, E., Winslow, R. L. & O’rourke, B. 2003. An integrated model of cardiac mitochondrial energy metabolism and calcium dynamics. Biophysical journal, 84, 2734–2755.

Dash, R. K., Li, Y., Kim, J., Beard, D. A., Saidel, G. M. & Cabrera, M. E. 2008. Metabolic dynamics in skeletal muscle during acute reduction in blood flow and oxygen supply to mitochondria: in-silico studies using a multi-scale, top-down integrated model. PloS one, 3, e3168.

Divakaruni, A. S. & Brand, M. D. 2011. The regulation and physiology of mitochondrial proton leak. Physiology (Bethesda), 26, 192–205.

Dong, Y., Digman, M. A. & Brewer, G. J. 2019. Age- and AD-related redox state of NADH in subcellular compartments by fluorescence lifetime imaging microscopy. Geroscience, 41, 51–67.

Filograna, R., Gerlach, J., Choi, H. N., Rigoni, G., Barbaro, M., Oscarson, M., Lee, S., Tiklova, K., Ringner, M., Koolmeister, C., Wibom, R., Riggare, S., Nennesmo, I., Perlmann, T., Wredenberg, A., Wedell, A., Motori, E., Svenningsson, P. & Larsson, N. G. 2024. PARKIN is not required to sustain OXPHOS function in adult mammalian tissues. NPJ Parkinsons Dis, 10, 93.

Gautier, C. A., Kitada, T. & Shen, J. 2008. Loss of PINK1 causes mitochondrial functional defects and increased sensitivity to oxidative stress. Proc Natl Acad Sci U S A, 105, 11364–9.

Ge, P., Dawson, V. L. & Dawson, T. M. 2020. PINK1 and Parkin mitochondrial quality control: a source of regional vulnerability in Parkinson’s disease. Mol Neurodegener, 15, 20.

Giguere, N., Pacelli, C., Saumure, C., Bourque, M. J., Matheoud, D., Levesque, D., Slack, R. S., Park, D. S. & Trudeau, L. E. 2018. Comparative analysis of Parkinson’s disease-associated genes in mice reveals altered survival and bioenergetics of Parkin-deficient dopamine neurons. J Biol Chem, 293, 9580–9593.

Haas, R. H., Nasirian, F., Nakano, K., Ward, D., Pay, M., Hill, R. & Shults, C. W. 1995. Low platelet mitochondrial complex I and complex II/III activity in early untreated Parkinson’s disease. Ann Neurol, 37, 714–22.

Hoepken, H. H., Gispert, S., Morales, B., Wingerter, O., Del Turco, D., Mulsch, A., Nussbaum, R. L., Muller, K., Drose, S., Brandt, U., Deller, T., Wirth, B., Kudin, A. P., Kunz, W. S. & Auburger, G. 2007. Mitochondrial dysfunction, peroxidation damage and changes in glutathione metabolism in PARK6. Neurobiol Dis, 25, 401–11.

Huber, H. J., Connolly, N. M., Dussmann, H. & Prehn, J. H. 2012. A structured approach to the study of metabolic control principles in intact and impaired mitochondria. Mol Biosyst, 8, 828–42.

Huber, H. J., Dussmann, H., Kilbride, S. M., Rehm, M. & Prehn, J. H. 2011. Glucose metabolism determines resistance of cancer cells to bioenergetic crisis after cytochrome-c release. Mol Syst Biol, 7, 470.

Jafri, M. S. & Kumar, R. 2014. Modeling mitochondrial function and its role in disease. Prog Mol Biol Transl Sci, 123, 103–25.

Joselin, A. P., Hewitt, S. J., Callaghan, S. M., Kim, R. H., Chung, Y. H., Mak, T. W., Shen, J., Slack, R. S. & Park, D. S. 2012. ROS-dependent regulation of Parkin and DJ-1 localization during oxidative stress in neurons. Hum Mol Genet, 21, 4888–903.

Kann, O. & Kovacs, R. 2007. Mitochondria and neuronal activity. Am J Physiol Cell Physiol, 292, C641–57.

Kitada, T., Pisani, A., Porter, D. R., Yamaguchi, H., Tscherter, A., Martella, G., Bonsi, P., Zhang, C., Pothos, E. N. & Shen, J. 2007. Impaired dopamine release and synaptic plasticity in the striatum of PINK1-deficient mice. Proc Natl Acad Sci U S A, 104, 11441–6.

Korge, P., Calmettes, G. & Weiss, J. N. 2015. Increased reactive oxygen species production during reductive stress: The roles of mitochondrial glutathione and thioredoxin reductases. Biochim Biophys Acta, 1847, 514–25.

Langston, J. W., Ballard, P., Tetrud, J. W. & Irwin, I. 1983. Chronic Parkinsonism in humans due to a product of meperidine-analog synthesis. Science, 219, 979–80.

Lin, M. T. & Beal, M. F. 2006. Mitochondrial dysfunction and oxidative stress in neurodegenerative diseases. Nature, 443, 787–95.

Liu, W., Acin-Perez, R., Geghman, K. D., Manfredi, G., Lu, B. & Li, C. 2011. Pink1 regulates the oxidative phosphorylation machinery via mitochondrial fission. Proc Natl Acad Sci U S A, 108, 12920–4.

Magnus, G. & Keizer, J. 1997. Minimal model of beta-cell mitochondrial Ca2+ handling. American Journal of Physiology-Cell Physiology, 273, C717–C733.

Malpartida, A. B., Williamson, M., Narendra, D. P., Wade-Martins, R. & Ryan, B. J. 2021. Mitochondrial Dysfunction and Mitophagy in Parkinson’s Disease: From Mechanism to Therapy. Trends Biochem Sci, 46, 329–343.

Morais, V. A., Haddad, D., Craessaerts, K., De Bock, P. J., Swerts, J., Vilain, S., Aerts, L., Overbergh, L., Grunewald, A., Seibler, P., Klein, C., Gevaert, K., Verstreken, P. & De Strooper, B. 2014. PINK1 loss-of-function mutations affect mitochondrial complex I activity via NdufA10 ubiquinone uncoupling. Science, 344, 203–7.

Moran, M., Moreno-Lastres, D., Marin-Buera, L., Arenas, J., Martin, M. A. & Ugalde, C. 2012. Mitochondrial respiratory chain dysfunction: implications in neurodegeneration. Free Radic Biol Med, 53, 595–609.

Muddapu, V. R. & Chakravarthy, V. S. 2021. Influence of energy deficiency on the subcellular processes of Substantia Nigra Pars Compacta cell for understanding Parkinsonian neurodegeneration. Sci Rep, 11, 1754.

Nicholls, D. G. 2013. Bioenergetics, Academic press.

Nicholls, D. G. & Budd, S. L. 2000. Mitochondria and neuronal survival. Physiol Rev, 80, 315–60.

Nicklas, W. J., Vyas, I. & Heikkila, R. E. 1985. Inhibition of NADH-linked oxidation in brain mitochondria by 1-methyl-4-phenyl-pyridine, a metabolite of the neurotoxin, 1-methyl-4-phenyl-1,2,5,6-tetrahydropyridine. Life Sci, 36, 2503–8.

Park, J. H., Burgess, J. D., Faroqi, A. H., Demeo, N. N., Fiesel, F. C., Springer, W., Delenclos, M. & Mclean, P. J. 2020. Alpha-synuclein-induced mitochondrial dysfunction is mediated via a sirtuin 3-dependent pathway. Mol Neurodegener, 15, 5.

Parker, W. D., JR., Parks, J. K. & Swerdlow, R. H. 2008. Complex I deficiency in Parkinson’s disease frontal cortex. Brain Res, 1189, 215–8.

Patro, S., Ratna, S., Yamamoto, H. A., Ebenezer, A. T., Ferguson, D. S., Kaur, A., Mcintyre, B. C., Snow, R. & Solesio, M. E. 2021. ATP Synthase and Mitochondrial Bioenergetics Dysfunction in Alzheimer’s Disease. Int J Mol Sci, 22.

Pauwels, P. J., Opperdoes, F. R. & Trouet, A. 1985. Effects of antimycin, glucose deprivation, and serum on cultures of neurons, astrocytes, and neuroblastoma cells. J Neurochem, 44, 143–8.

Pedregosa, F., Varoquaux, G., Gramfort, A., Michel, V., Thirion, B., Grisel, O., Blondel, M., Prettenhofer, P., Weiss, R. & Dubourg, V. 2011. Scikit-learn: Machine learning in Python. the Journal of machine Learning research, 12, 2825–2830.

Pilsl, A. & Winklhofer, K. F. 2012. Parkin, PINK1 and mitochondrial integrity: emerging concepts of mitochondrial dysfunction in Parkinson’s disease. Acta Neuropathol, 123, 173–88.

Rossi, A., Rigotto, G., Valente, G., Giorgio, V., Basso, E., Filadi, R. & Pizzo, P. 2020. Defective Mitochondrial Pyruvate Flux Affects Cell Bioenergetics in Alzheimer’s Disease-Related Models. Cell Rep, 30, 2332–2348 e10.

Rovini, A., Heslop, K., Hunt, E. G., Morris, M. E., Fang, D., Gooz, M., Gerencser, A. A. & Maldonado, E. N. 2021. Quantitative analysis of mitochondrial membrane potential heterogeneity in unsynchronized and synchronized cancer cells. FASEB J, 35, e21148.

Sadri, S., Zhang, X., Audi, S. H., Cowley, A. W., JR. & Dash, R. K. 2023. Computational Modeling of Substrate-Dependent Mitochondrial Respiration and Bioenergetics in the Heart and Kidney Cortex and Outer Medulla. Function (Oxf), 4, zqad038.

Saltelli, A., Tarantola, S., Campolongo, F. & Ratto, M. 2004. Sensitivity analysis in practice: a guide to assessing scientific models. Chichester, England.

Scaduto, R. C. & Grotyohann, L. W. 1999. Measurement of mitochondrial membrane potential using fluorescent rhodamine derivatives. Biophysical journal, 76, 469–477.

Schapira, A. H. 2008. Mitochondria in the aetiology and pathogenesis of Parkinson’s disease. Lancet Neurol, 7, 97–109.

Schapira, A. H., Mann, V. M., Cooper, J. M., Dexter, D., Daniel, S. E., Jenner, P., Clark, J. B. & Marsden, C. D. 1990. Anatomic and disease specificity of NADH CoQ1 reductase (complex I) deficiency in Parkinson’s disease. J Neurochem, 55, 2142–5.

Schmidt, C. A., Fisher-Wellman, K. H. & Neufer, P. D. 2021. From OCR and ECAR to energy: Perspectives on the design and interpretation of bioenergetics studies. J Biol Chem, 297, 101140.

Sipos, I., Tretter, L. & Adam-Vizi, V. 2003. Quantitative relationship between inhibition of respiratory complexes and formation of reactive oxygen species in isolated nerve terminals. J Neurochem, 84, 112–8.

Theurey, P., Connolly, N. M., Fortunati, I., Basso, E., Lauwen, S., Ferrante, C., Moreira Pinho, C., Joselin, A., Gioran, A. & Bano, D. 2019. Systems biology identifies preserved integrity but impaired metabolism of mitochondria due to a glycolytic defect in Alzheimer’s disease neurons. Aging cell, 18, e12924.

Wang, W., Zhao, F., Ma, X., Perry, G. & Zhu, X. 2020. Mitochondria dysfunction in the pathogenesis of Alzheimer’s disease: recent advances. Mol Neurodegener, 15, 30.

Ward, M. W., Rego, A. C., Frenguelli, B. G. & Nicholls, D. G. 2000. Mitochondrial membrane potential and glutamate excitotoxicity in cultured cerebellar granule cells. J Neurosci, 20, 7208–19.

Waskom, M. L. 2021. Seaborn: statistical data visualization. Journal of Open Source Software, 6, 3021.

Wu, F., Yang, F., Vinnakota, K. C. & Beard, D. A. 2007. Computer modeling of mitochondrial tricarboxylic acid cycle, oxidative phosphorylation, metabolite transport, and electrophysiology. Journal of Biological Chemistry, 282, 24525–24537.

Yin, F., Boveris, A. & Cadenas, E. 2014. Mitochondrial energy metabolism and redox signaling in brain aging and neurodegeneration. Antioxid Redox Signal, 20, 353–71.

