## Supplementary Methods for "Integrating simulated and experimental data to identify mitochondrial bioenergetic defects in Parkinson’s Disease models"

**Supplementary Methods – Tables adapted from (Theurey et al., 2019)**

**Table S1: Modelled state variables, their initial concentrations (following steady-state calculations) and literature references where available.**

| **State Variable** | **Description** | **Initial Conc.** | **Unit** | **Reference** |
| --- | --- | --- | --- | --- |
| H_x_ | Proton (H^+^; matrix) | 1.8x10^-8^ | M | pH = 7.7; (Bolshakov et al., 2008) |
| K_x_ | Potassium (K^+^; matrix) | 55 | mM |  |
| Mg_x_ | Magnesium (Mg^2+^; matrix) | 400 | μM |  |
| NADH_x_ | NADH (matrix) | 350 | μM | (Wei et al., 2011) |
| QH_2_ | Ubiquinol (matrix) | 40 | μM |  |
| C_red_ | Reduced cytochrome-*c* | 180 | μM |  |
| ATP_x_ | Total ATP (matrix) | 1.1 | mM | (Yoshida et al., 2016) |
| ADP_x_ | Total ADP (matrix) | 1.5 | mM |  |
| ATP_mx_ | ATP bound to Mg^2+^ (matrix) | 730 | μM |  |
| ADP_mx_ | ADP bound to Mg^2+^ (matrix) | 830 | μM |  |
| Pi_x_ | Inorganic phosphate (matrix) | 16 | mM |  |
| ATP_i_ | Total ATP (IMS) | 2.3 | mM |  |
| ADP_i_ | Total ADP (IMS) | 290 | μM |  |
| AMP_i_ | Total AMP (IMS) | 0 | M |  |
| ATP_mi_ | ATP bound to Mg^2+^ (IMS) | 2.3 | mM |  |
| ADP_mi_ | ADP bound to Mg^2+^ (IMS) | 290 | μM |  |
| Pi_i_ | Inorganic phosphate (IMS) | 20 | mM |  |
| dPsi | Mitochondrial membrane potential (ψΔ_m_) | 149 | mV | (Gerencser et al., 2012, Ward et al., 2000, Nicholls and Budd, 2000) |
| H_i_ | Hydrogen (H^+^, protons; IMS) | 3.9x10^-8^ | M |  |
| ATP_c_ | ATP (cytosol) | 2.3 | mM | (Nelson et al., 2008) |
| ADP_c_ | ADP (cytosol) | 290 | μM | ATP_c_/ADP_c_ ≈ 10; (Hardie and Hawley, 2001, Katsura et al., 1993) |
| H_2_O_2_ | H_2_O_2_ (cytosol) | 10^-8^ | M |  |

*Suffixes: _x_, mitochondrial matrix; _i_, mitochondrial intermembrane space (IMS); _c_, cytosolic; _m_, Mg^2+^ bound; _red_, reduced; _tot_, total*

**Table S2: Ordinary differential equations (ODEs) describing the state variables listed in Table S1. Fluxes (J_*) and parameters contributing to the ODEs are described in Tables S3 and S4. Detailed explanation of these ODEs can be found in (Beard, 2005, Huber et al., 2011, Chenna et al., 2021).**

| **State Variable** | **Ordinary Differential Equation** | **Note** |
| --- | --- | --- |
| H_x_ | $\frac{x_{buff} * H_{x} * (+1*J_{DH} - (4+1)*J_{C1} - (4-2)*J_{C3} - (2+2)*J_{C4} + (n_{A}-1)*J_{F1} + 2*J_{Pi1} + J_{Hle} - J_{KH})}{W_{x}}$ |  |
| K_x_ | $\frac{J_{KH} + J_{K}}{W_{x}}$ |  |
| Mg_x_ | $\frac{{-J_{MgATP}}_{x}{-J_{MgADP}}_{x}}{W_{x}}$ |  |
| NADH_x_ | $\frac{J_{DH} - J_{C1}}{W_{x}}$ |  |
| QH_2_ | $\frac{J_{C1} - J_{C3}}{W_{x}}$ | 1 |
| C_red_ | $\frac{2J_{C3} - 2J_{C4}}{W_{i}}$ |  |
| ATP_x_ | $\frac{J_{F1} - J_{ANT}}{W_{x}}$ |  |
| ADP_x_ | $\frac{-J_{F1} + J_{ANT}}{W_{x}}$ |  |
| ATP_mx_ | $\frac{{J_{MgATP}}_{x}}{W_{x}}$ |  |
| ADP_mx_ | $\frac{{J_{MgADP}}_{x}}{W_{x}}$ |  |
| Pi_x_ | $\frac{-J_{F1} + J_{Pi1}}{W_{x}}$ |  |
| ATP_i_ | $\frac{J_{ATP} +J_{ANT} + {J_{AK}}_{i}}{W_{i}}$ |  |
| ADP_i_ | $\frac{J_{ADP} -J_{ANT} - 2{J_{AK}}_{i}}{W_{i}}$ |  |
| AMP_i_ | $\frac{J_{AMP} + {J_{AK}}_{i}}{W_{i}}$ |  |
| ATP_mi_ | $\frac{{J_{MgATP}}_{i}}{W_{i}}$ |  |
| ADP_mi_ | $\frac{{J_{MgADP}}_{i}}{W_{i}}$ |  |
| Pi_i_ | $\frac{-J_{Pi1} + J_{Pi2}}{W_{i}}$ |  |
| dPsi | $\frac{4J_{C1} + 2J_{C3} + 4J_{C4} - n_{A}*J_{F1} - J_{\mathrm{ANT}} - J_{\mathrm{Hle}} - J_{K}}{C_{\mathrm{IM}}}$ |  |
| H_i_ | $\frac{-J_{DH} + (4+1)*J_{C1} + (4-2)*J_{C3} + (2+2)*J_{C4} - (n_{A}-1)*J_{F1} - 2*J_{Pi1} - J_{Hle} + J_{KH} + J_{Ht})}{W_{i}}$ |  |
| ATP_c_ | $\frac{-J_{ATP} - J_{ATPK}}{W_{c}}$ |  |
| ADP_c_ | $\frac{-J_{ADP} + J_{ATPK}}{W_{c}}$ |  |
| H_2_O_2_ | $\frac{V\_H_{2}O_{2}\_If+V\_H_{2}O_{2}\_Qo -k\_H_{2}O_{2}*H_{2}O_{2}}{W_{c}}$ |  |

*Suffixes: _x_, mitochondrial matrix; _i_, mitochondrial intermembrane space (IMS); _c_, cytosolic; _m_, Mg^2+^ bound; _red_, reduced; _tot_, total*

*^1^As the inner mitochondrial membrane is not explicitly modelled as a separate compartment, the model describes the binding of QH_2_ to the respiratory complexes without specifying the precise location of this event. QH_2_ concentration is simulated as the concentration in the mitochondrial matrix*

**Table S3: Fluxes contributing to the ODEs described in Table S2, as described previously (Beard, 2005, Chenna et al., 2021). Parameters contributing to these fluxes are described in Table S4.**

| **Flux** | **Description** | | **Equation** | | | **Unit** |
| --- | --- | --- | --- | --- | --- | --- |
| Membrane proton-motive forces and respiration fluxes | | | | | |  |
| dG_H_ | Protomotive force | | $F*dPsi+1*RT*\log\left( \frac{H_{i}}{H_{x}} \right)$ | | |  |
| dG_C1op_ | Gibb’s energy complex I | | $dG_{C1o}-1*RT*log(\frac{H_{x}}{1e^{-7}})$ | | |  |
| dG_C3op_ | Gibb’s energy complex III | | $dG_{C3o}+2*RT*log(\frac{H_{x}}{1e^{-7}})$ | | |  |
| dG_C4op_ | Gibb’s energy complex IV | | $dG_{C4o}-2*RT*log(\frac{H_{x}}{1e^{-7}})$ | | |  |
| dG_F1op_ | Gibb’s energy F_1_F_o_ ATP synthase | | $dG_{F1o}-1*RT*log(\frac{H_{x}}{1e^{-7}})$ | | |  |
| J_DH_ | Mitochondrial dehydrogenase flux (input NADH flux) | | $x_{DH}*\left( r_{DH}*NAD_{x}-NADH_{x} \right)*\frac{1+{Pi_{x}}/{k_{Pi1}}}{1+{Pi_{x}}/{k_{Pi2}}}$ | | |  |
| J_C1_ | Flux through complex I | | $x_{c1}*\left( \exp\left( \frac{-\left( dG_{C1op}+4dG_{H} \right)}{RT} \right)*NA{DH}_{x}*Q-NAD_{x}*QH_{2} \right)$ | | | mol s^-1^ (l mito)^-1^ |
| J_C3_ | Flux through complex III | | $x_{c3}*\frac{1+{Pi_{x}}/{k_{Pi3}}}{1+{Pi_{x}}/{k_{Pi4}}} *\exp\left( \frac{-\left( dG_{C3op}+4dG_{H}-2F*dPsi \right)}{2RT} \right)*C_{ox}*\surd QH_{2}-C_{red}*\surd Q$ | | | mol s^-1^ (l mito)^-1^ |
| J_C4_ | Flux through complex IV | | $x_{c4}*\frac{O_{2}}{O_{2}+k_{O2}} *\frac{C_{red}}{C_{tot}}\exp\left( \frac{-\left( dG_{C4op}+2dG_{H} \right)}{2RT} \right)*C_{red}*O_{2}^{0.25}-C_{ox}*exp(F*\frac{dPsi}{RT})$ | | | mol s^-1^ (l mito)^-1^ |
| J_F1_ | Flux through F_1_F_o_ ATP synthase | | $x_{F1}*\left( \exp\left( \frac{-\left( dG_{F1op}-n_{A}*dG_{H} \right)}{RT} \right)*\frac{K_{DD}}{K_{DT}}*ADP_{mx}*Pi_{x}-ATP_{mx} \right)$ | | | mol s^-1^ (l mito)^-1^ |
| ATP transferase | | | | | |  |
| J_ANT_ | Adenosine nucleotide transferase | | $x_{ANT}*\left( \frac{ADP_{fi}}{ADP_{fi}+ATP_{fi}*\exp\left( -F*\frac{Psi_{i}}{RT} \right)}-\frac{ADP_{fx}}{ADP_{fx}+ATP_{fx}*\exp\left( -F*\frac{Psi_{x}}{RT} \right)} \right)* \frac{{ADP}_{fi}}{ADP_{fi}+k_{mADP}}$ | | | mol s^-1^ (l mito)^-1^ |
| Ionic Fluxes |  | |  | | |  |
| H_2_Pi_i_ | Pi H^+^ binding (IMS) | | $Pi_{i}*H_{i}*\left( H_{i}+k_{dHPi} \right)$ | | |  |
| H_2_Pi_x_ | Pi H^+^ binding (matrix) | | $Pi_{x}*\frac{H_{x}}{\left( H_{x}+k_{dHPi} \right)}$ | | |  |
| J_Pi1_ | Pi/H^+^ exchanger | | $x_{Pi1}*\frac{H_{x}*H_{2}{Pi}_{i}-H_{i}*H_{2}{Pi}_{x}}{H_{2}Pi_{i}+k_{PiH}}$ | | | mol s^-1^ (l mito)^-1^ |
| J_Hle_ | Proton (H^+^) leaks | | $x_{Hle}*dPsi*\frac{H_{i}*exp(F*\frac{dPsi}{RT})-H_{x}}{\exp\left( F*\frac{dPsi}{RT} \right)-1}$ | | | mol s^-1^ (l mito)^-1^ |
| J_KH_ | K^+^/ H^+^ exchanger | | $x_{KH}*\left( K_{i}*H_{x}-K_{x}*H_{i} \right)$ | | | mol s^-1^ (l mito)^-1^ |
| J_K_ | K^+^ uniport | | $x_{K}*dPsi*\frac{K_{i}*\exp\left( F*\frac{dPsi}{RT} \right)-K_{x}}{\exp\left( F*\frac{dPsi}{RT} \right)-1}$ | | |  |
| J_AKi_ | Mitochondrial adenylate kinase | | $x_{AK}*\left( K_{AK}*ADP_{i}*ADP_{i}-AMP_{i}*ATP_{i} \right)$ | | | mol s^-1^ (l mito)^-1^ |
| J_AMP_ | AMP MOM permeability | | $gamma*x_{A}*\left( AMP_{e}-AMP_{i} \right)$ | | | mol s^-1^ (l mito)^-1^ |
| J_ADP_ | ADP MOM permeability | | $gamma*x_{A}*\left( ADP_{e}-ADP_{i} \right)$ | | | mol s^-1^ (l mito)^-1^ |
| J_ATP_ | ATP MOM permeability | | $gamma*x_{A}*\left( ATP_{e}-ATP_{i} \right)$ | | | mol s^-1^ (l mito)^-1^ |
| J_Pi2_ | Phosphate MOM permeability | | $gamma*x_{Pi2}*\left( {Pi}_{e}-{Pi}_{i} \right)$ | | |  |
| J_Ht_ | Proton MOM permeability | | $gamma*x_{Ht}*\left( H_{e}-H_{i} \right)$ | | |  |
| J_MgATPx_ | Mg^2+^ binding to ATP_x_ | | $x_{MgA}*\left( ATP_{fx}*Mg_{x}-K_{DT}*ATP_{mx} \right)$ | | |  |
| J_MgADPx_ | Mg^2+^ binding to ADP_x_ | | $x_{MgA}*\left( ADP_{fx}*Mg_{x}-K_{DD}*ADP_{mx} \right)$ | | |  |
| J_MgATPi_ | Mg^2+^ binding to ATPi | | $x_{MgA}*\left( ATP_{fi}*Mg_{i}-K_{DT}*ATP_{mi} \right)$ | | |  |
| J_MgADPi_ | Mg^2+^ binding to ADPi | | $x_{MgA}*\left( ADP_{fi}*Mg_{i}-K_{DD}*ADP_{mi} \right)$ | | |  |
| Cytosolic Energy balance to model intact cells | | | | | |  |
| J_ATPK_ | Cytosolic ATP processes | | $x_{ATPK}*\left( ATP_{e}-K_{ADTP\_dyn}*ADP_{e} \right)$ | | |  |
| Reactive oxygen species (H_2_O_2_) metabolism (Chenna et al., 2021) | | | | | | |
| V_$H_{2}O_{2}$_If | | H_2_O_2_ from site I_f_ | | ${(a}_{IF}.e^{\left( m_{IF}.\frac{{NADH}_{x}}{{NAD}_{tot}} \right)}-c_{IF}).(d_{IF}+e_{IF}\left( 1-\frac{1}{1+e^{\left( \frac{{\Delta\varphi}_{m}-t_{IF}}{k_{If}} \right)}} \right)).x_{IF}$ | mol s^-1^ (l mito)^-1^ | |
| V_$H_{2}O_{2}$_Qo | | H_2_O_2_ from site Q_o_ | | ${(a}_{QO}.e^{\left( m_{QO}.\frac{{QH}_{2}}{Q_{tot}} \right)}-c_{QO}).x_{QO}$ | mol s^-1^ (l mito)^-1^ | |

*Suffixes: x, mitochondrial matrix; i, mitochondrial intermembrane space (IMS); c, cytosolic; e, cytosolic (external); m, Mg^2+^ bound; red, reduced; tot, total*

*Abbreviations: MOM, mitochondrial outer membrane; Pi, phosphate*

**Table S4: Model parameters, their calibrated values, and literature references where available**

| **Symbol** | **Description** | **Value** | **Unit** | |  | **Reference** |
| --- | --- | --- | --- | --- | --- | --- |
| NAD_tot_ | Total NAD(H) concentration (NAD^+^+NADH) | 726x10^-6^ | M | | c) | (Wei et al., 2011, Cloutier et al., 2009) |
| C_tot_ | Total cytochrome-*c* concentration (C_red_+C_ox_) | 2.7x10^-3^ | M | | b) | (Beard, 2005) |
| Q_tot_ | Total ubiquinol/ubiquinone concentration (Q+QH_2_) | 1.4x10^-3^ | M | | b) | (Dash and Beard, 2008) |
| ADTP_tot_ | Total cytosolic adenosine phosphates | 2.6x10^-3^ | M | | a) | (Cloutier et al., 2009, DiNuzzo et al., 2010) |
| O_2_ | Oxygen concentration | 26x10^-6^ | M | | a) | (Aubert and Costalat, 2005, Dmitriev et al., 2015, Murphy, 2009) |
| Dehydrogenase flux input function | |  |  | |  |  |
| k_Pi1_ | Dehydrogenase flux input (phosphate dependency) | 0.13x10^-3^ | M | | b) | (Beard, 2005) |
| k_Pi2_ | Dehydrogenase flux input (phosphate dependency) | 0.68x10^-3^ | M | | b) | (Beard, 2005) |
| x_DH_ | Dehydrogenase activity (flux activity) | 59 x10^-3^ | mol s^-1^ M^-1^ (l mito vol)^-1^ | | c) | (Dash and Beard, 2008) |
| r_DH_ | NADH/NAD^+^ equilibrium constant | 4.3 | Unitless | | c) | (Beard, 2005) |
| Complex I | |  |  | |  |  |
| x_C1_ | Complex I activity | 1020 | mol s^-1^ M^-2^ (l mito vol)^-1^ | | b) | (Huber et al., 2012) |
| Complex III | |  |  | |  |  |
| x_C3_ | Complex III activity | 224 x10^-3^ | mol s^-1^ M^-3/2^ (l mito vol)^-1^ | | b) | (Huber et al., 2012) |
| k_Pi3_ | Pi dependency parameter 1 | 0.19x10^-3^ | M | | b) | (Beard, 2005) |
| k_Pi4_ | Pi dependency parameter 2 | 25x10^-3^ | M | | b) | (Beard, 2005) |
| Complex IV | |  |  | |  |  |
| x_C4_ | Complex IV activity | 0.32 x10^-3^ | mol s^-1^ M^-1^ (l mito vol)^-1^ | | b) | (Huber et al., 2012) |
| k_O2_ | Saturation constant for oxygen consumption | 0.12 x10^-3^ | M | | b) | (Dash and Beard, 2008) |
| ATP synthase | |  |  | |  |  |
| x_F1_ | F_1_F_o_ ATP synthase activity | 6829 | mol s^-1^ M^-1^ (l mito vol)^-1^ | | b) | (Huber et al., 2012) |
| Mg-binding to ATP/ADP | |  |  | |  |  |
| K_DT_ | Mg^2+^/ATP binding constant | 192x10^-6^ | M | | c) |  |
| K_DD_ | Mg^2+^/ADP binding constant | 347 x10^-6^ | M | | b) | (Dash and Beard, 2008) |
| x_MgA_ | Mg^2+^ binding activity | 1 x10^6^ |  | | b) | (Beard, 2005) |
| Cytosolic ATP production and consumption | |  |  | |  |  |
| x_ATPK_ | Cytosolic ATP production & consumption activity | 0.5 | min^-1^ | | c) |  |
| K_ADTP_dyn_ | ATP consumption equilibrium constant | 3.4 | Unitless | | c) |  |
| Adenosine transferase | |  |  | |  |  |
| x_ANT_ | ANT activity | 2 x10^-3^ | mol s^-1^ (l mito vol)^-1^ | | b) | (Huber et al., 2011) |
| k_mADP_ | ANT parameter 1 | 3.5 x10^-6^ | M | | b) | (Dash and Beard, 2008) |
| Proton leaks | |  |  | |  |  |
| x_Hle_ | Proton leak activity | 150 | mol s^-1^ M^-1^ mV^-1^ (l mito vol)^-1^ | | b) | (Huber et al., 2011, Dash and Beard, 2008) |
| OM transporters | |  |  | |  |  |
| x_Ht_ | MOM permeability to protons | 2000 | μm s^-1^ | | a) | ^2^ |
| gamma | MOM area per unit (mito) volume | 5.99 | μm^-1^ | | b) | (Beard, 2005) |
| x_A_ | MOM permeability to nucleotides | 85 | μm s^-1^ (l mito)^-1^ | | b) | (Dash and Beard, 2008) |
| x_Pi2_ | MOM permeability to phosphate | 327 | μm s^-1^ (l mito)^-1^ | | b) | (Dash and Beard, 2008) |
| Phosphate-hydrogen co-transport | |  |  | |  |  |
| k_dHPi_ | H^+^/Pi co-transport binding constant | 10^-6.75^ | M | | b) | (Dash and Beard, 2008) |
| k_PiH_ | H^+^/Pi co-transport dissociation constant | 0.45 x10^-3^ | M | | b) | (Beard, 2005) |
| x_Pi1_ | H^+^/Pi co-transport activity | 385 x10^3^ | mol s^-1^ M^-1^ (l mito vol)^-1^ | | b) | (Dash and Beard, 2008) |
| Potassium-hydrogen anti-port | |  |  | |  |  |
| x_KH_ | K^+^/H^+^ antiporter activity | 29.8 x10^6^ | mol s^-1^ M^-2^ (l mito vol)^-1^ | | b) | (Beard, 2005) |
| Membrane buffer and capacitance | |  |  | |  |  |
| x_buff_ | Inner matrix H^+^ buffer capacity | 100 | M^-1^ | | b) | (Beard, 2005) |
| C_IM_ | Mitochondrial inner membrane capacitance | 6.8 x10^-6^ | mol (l mito vol)^-1^ mV^-1^ | | b) | (Dash and Beard, 2008) |
| Thermodynamic parameters | |  |  | |  |  |
| F | Faraday's constant | 0.096 | kJ mol^-1^ mV^-1^ | |  |  |
| R | Universal gas constant | 8314x10^-6^ | kJ mol^-1^ K^-1^ | |  |  |
| T | Temperature | 310.15 | K | |  |  |
| Gibbs free energy | |  |  | |  |  |
| dG_C10_ | Gibbs free energy for complex I/II reaction at pH 7 | -69.37 | kJ mol^-1^ | | b) | (Dash and Beard, 2008) |
| dG_C30_ | Gibbs free energy for complex III reaction | -32.53 | kJ mol^-1^ | | b) | (Dash and Beard, 2008) |
| dG_C40_ | Gibbs free energy for complex IV reaction | -122.94 | kJ mol^-1^ | | b) | (Dash and Beard, 2008) |
| dG_F10_ | Gibbs free energy for ATP synthase reaction | 36.03 | kJ mol^-1^ | | b) | (Dash and Beard, 2008) |
| n_A_ | Number of protons pumped by ATP synthase | 3 | Unitless | | b) | (Dash and Beard, 2008) |
| Cytosolic ion/nucleotide concentrations and pH | |  |  | |  |  |
| pH_e_ | pH (cytosol) | 7.4 | Unitless | | a) | (Bolshakov et al., 2008) |
| H_e_ | H^+^ concentration (cytosol) | 10^-pHe^ | M | | b) | (Huber et al., 2011) |
| K_ei_ | K^+^ concentration (cytosol) | 120x10^-3^ | M | | a) | (Liu et al., 2003) |
| Mg_tot_ | Mg^2+^ concentration (cytosol) | 20x10^-3^ | M | | a) | (Kubota et al., 2005) |
| Pi_e_ | Phosphate (Pi) concentration (cytosol) | 20x10^-3^ | M | | b) | (Huber et al., 2011) |
| state_fact | Maintains steady-state ATP:ADP at 10:1 | 10/11 | Unitless | | b) | (Huber et al., 2011) |
| ATP_e | Steady-state ATP concentration (cytosol) | state_fact*ADTP__tot_ | M | |  |  |
| ADP_e_ | Steady-state ADP concentration (cytosol) | ADTP__tot_-ATP__e_ | M | |  |  |
| AMP_e_ | AMP concentration (cytosol) | 0 | M | |  |  |
| Mitochondrial volume/fraction | |  |  | |  |  |
| V_mito_ | Mitochondrial Fraction | 0.06 | % | | a) | (Ward et al., 2007) |
| W_c_ | Volume fraction: cytosol/mitochondria | 1/V__mito_ |  | |  |  |
| W_m_ | Mitochondrial water space | 0.7 | ml water/ml mito | | b) | (Beard, 2005) |
| W_x_ | Volume fraction: matrix/mitochondria | 0.9*W_m_ | ml water/ml mito | | b) | (Beard, 2005) |
| W_i_ | Volume fraction: IMS/mitochondria | 0.1*W_m_ | ml water/ml mito | | b) | (Beard, 2005) |
| Potassium uniport and adenylate kinase (neglected) | | |  | |  | (Beard, 2005, Huber et al., 2011) |
| x_K_ | Passive potassium transporter activity | 0 |  | |  |  |
| x_AK_ | Adenylate kinase activity | 0 |  | |  |  |
| K_AK_ |  | 0 |  | |  |  |
| Moiety conservations | |  |  | |  |  |
| NAD_x_ | Total NAD - reduced NADH (matrix) | NAD_tot_ - NADH_x_ |  | |  |  |
| Q | Total ubiquinol - reduced ubiquinol | Q_tot_ - QH_2_ |  | |  |  |
| C_ox_ | IMS Total cyt *c* - IMS reduced cyt *c* | C_tot_ - C_red_ |  | |  |  |
| ATP_fx_ | Free ATP_x_ | ATP_x_ - ATP_mx_ |  | |  |  |
| ADP_fx_ | Free ADP_x_ | ADP_x_ - ADP_mx_ |  | |  |  |
| ATP_fi_ | Free ATP_i_ | ATP_i_ - ATP_mi_ |  | |  |  |
| ADP_fi_ | Free ADP_i_ | ADP_i_ - ADP_mi_ |  | |  |  |
| ADP_me_ | Mg-bound ADP |  | | $\frac{\left( K_{DD}+ADP_{e}+Mg_{tot} \right)-\sqrt{\left( K_{DD}+ADP_{e}+Mg_{tot} \right)^{2}-4\left( Mg_{tot}*ADP_{e} \right)}}{2}$ | | |
| ADP_fe_ | Free cytosolic ADP | ADP_e_ - ADP_me_ |  | |  |  |
| Mg_e_ | Free Mg^2+^ concentration | Mg_tot_ - ADP_me_ |  | |  |  |
| Mg_i_ | Mg^2+^ (IMS)^1^ | Mg_e_ |  | |  |  |
| K_i_ | K^+^ (IMS)^1^ | K_ei_ |  | |  |  |
| Parameter for Adenine Nucleotide Transferase (ANT) | |  |  | |  |  |
| Psi_x_ |  | -0.65*dPsi |  | | b) | (Korzeniewski and Brown, 1998) |
| Psi_i_ |  | 0.35*dPsi |  | | b) | (Korzeniewski and Brown, 1998) |
| Reactive oxygen species metabolism (Chenna et al., 2021) | | | | | | |
| aIf | Complex I regression constant 1 | 0.142 | Unitless | | b) | (Chenna et al., 2021) |
| mIf | Complex I regression constant 2 | 2.7 | Unitless | | b) | (Chenna et al., 2021) |
| cIf | Complex I regression intercept | 1.003 | Unitless | | b) | (Chenna et al., 2021) |
| dIf | ΔΨ_m_-independent H_2_O_2_ production | 0.37 | Unitless | | b) | (Chenna et al., 2021) |
| eIf | ΔΨ_m_-dependent H_2_O_2_ production | 0.63 | Unitless | | b) | (Chenna et al., 2021) |
| tIf | Half-maximal voltage (V_1/2_; voltage at which ΔΨ_m_-dependent H_2_O_2_ production is half maximal) | 169 | mV | | b) | (Chenna et al., 2021) |
| tIf | Boltzmann slope factor | 1.22 | mV | | b) | (Chenna et al., 2021) |
| xIf | Complex I H_2_O_2_ scaling factor | 1.2e^−7^ | M.s^-1^ | |  | (Chenna et al., 2021) |
| aQo | Complex III regression constant 1 | 0.046176 | unitless | | b) | (Chenna et al., 2021) |
| mQo | Complex III regression constant 2 | 6.03612 | unitless | | b) | (Chenna et al., 2021) |
| cQo | Complex III regression intercept | 0.0796868 | unitless | | b) | (Chenna et al., 2021) |
| xQo | Complex III H_2_O_2_ scaling factor | 1e^−8^ | M.s^-1^ | | b) | (Chenna et al., 2021) |
| k_$H_{2}O_{2}$ | First order H_2_O_2_ scavenging constant | 120 | s^-1^ | | a) | (Starkov et al., 2002) |

*Suffixes: x, mitochondrial matrix; i, mitochondrial intermembrane space (IMS); c, cytosolic; e, cytosolic (external); m, Mg^2+^ bound; red, reduced; tot, total*

*Abbreviations: MOM, mitochondrial outer membrane; Pi, phosphate*

*^1^* K^+^ and Mg^2+^ ions rapidly equilibrate across OM, so IMS concentration assumed to be equal to cytosolic concentration

^2^ Proton permeability across MOM is assumed >> than permeability of nucleotides and phosphate

*a) Value set ~according to literature*

*b) Value retained from previous publications utilising model*

*c) Value adjusted during model calibration to in-house measurements in (Theurey et al., 2019)*

**Supplementary References**

AUBERT, A. & COSTALAT, R. 2005. Interaction between astrocytes and neurons studied using a mathematical model of compartmentalized energy metabolism. *J Cereb Blood Flow Metab,* 25**,** 1476-90.

BEARD, D. A. 2005. A biophysical model of the mitochondrial respiratory system and oxidative phosphorylation. *PLoS Comput Biol,* 1**,** e36.

BOLSHAKOV, A. P., MIKHAILOVA, M. M., SZABADKAI, G., PINELIS, V. G., BRUSTOVETSKY, N., RIZZUTO, R. & KHODOROV, B. I. 2008. Measurements of mitochondrial pH in cultured cortical neurons clarify contribution of mitochondrial pore to the mechanism of glutamate-induced delayed Ca2+ deregulation. *Cell Calcium,* 43**,** 602-14.

CHENNA, S., PREHN, J. H. & CONNOLLY, N. M. Phenomenological equations for electron transport chain-mediated reactive oxygen species metabolism. 2021 IEEE International Conference on Bioinformatics and Biomedicine (BIBM), 2021. IEEE, 653-658.

CLOUTIER, M., BOLGER, F. B., LOWRY, J. P. & WELLSTEAD, P. 2009. An integrative dynamic model of brain energy metabolism using in vivo neurochemical measurements. *J Comput Neurosci,* 27**,** 391-414.

DASH, R. K. & BEARD, D. A. 2008. Analysis of cardiac mitochondrial Na+-Ca2+ exchanger kinetics with a biophysical model of mitochondrial Ca2+ handling suggests a 3:1 stoichiometry. *J Physiol,* 586**,** 3267-85.

DINUZZO, M., MANGIA, S., MARAVIGLIA, B. & GIOVE, F. 2010. Changes in glucose uptake rather than lactate shuttle take center stage in subserving neuroenergetics: evidence from mathematical modeling. *J Cereb Blood Flow Metab,* 30**,** 586-602.

DMITRIEV, R. I., BORISOV, S. M., KONDRASHINA, A. V., PAKAN, J. M., ANILKUMAR, U., PREHN, J. H., ZHDANOV, A. V., MCDERMOTT, K. W., KLIMANT, I. & PAPKOVSKY, D. B. 2015. Imaging oxygen in neural cell and tissue models by means of anionic cell-permeable phosphorescent nanoparticles. *Cell Mol Life Sci,* 72**,** 367-81.

GERENCSER, A. A., CHINOPOULOS, C., BIRKET, M. J., JASTROCH, M., VITELLI, C., NICHOLLS, D. G. & BRAND, M. D. 2012. Quantitative measurement of mitochondrial membrane potential in cultured cells: calcium-induced de- and hyperpolarization of neuronal mitochondria. *J Physiol,* 590**,** 2845-71.

HARDIE, D. G. & HAWLEY, S. A. 2001. AMP-activated protein kinase: the energy charge hypothesis revisited. *Bioessays,* 23**,** 1112-9.

HUBER, H. J., CONNOLLY, N. M., DUSSMANN, H. & PREHN, J. H. 2012. A structured approach to the study of metabolic control principles in intact and impaired mitochondria. *Mol Biosyst,* 8**,** 828-42.

HUBER, H. J., DUSSMANN, H., KILBRIDE, S. M., REHM, M. & PREHN, J. H. 2011. Glucose metabolism determines resistance of cancer cells to bioenergetic crisis after cytochrome-c release. *Mol Syst Biol,* 7**,** 470.

KATSURA, K., RODRIGUEZ DE TURCO, E. B., FOLBERGROVA, J., BAZAN, N. G. & SIESJO, B. K. 1993. Coupling among energy failure, loss of ion homeostasis, and phospholipase A2 and C activation during ischemia. *J Neurochem,* 61**,** 1677-84.

KORZENIEWSKI, B. & BROWN, G. C. 1998. Quantification of the relative contribution of parallel pathways to signal transfer: application to cellular energy transduction. *Biophys Chem,* 75**,** 73-80.

KUBOTA, T., SHINDO, Y., TOKUNO, K., KOMATSU, H., OGAWA, H., KUDO, S., KITAMURA, Y., SUZUKI, K. & OKA, K. 2005. Mitochondria are intracellular magnesium stores: investigation by simultaneous fluorescent imagings in PC12 cells. *Biochim Biophys Acta,* 1744**,** 19-28.

LIU, D., SLEVIN, J. R., LU, C., CHAN, S. L., HANSSON, M., ELMER, E. & MATTSON, M. P. 2003. Involvement of mitochondrial K+ release and cellular efflux in ischemic and apoptotic neuronal death. *J Neurochem,* 86**,** 966-79.

MURPHY, M. P. 2009. How mitochondria produce reactive oxygen species. *Biochem J,* 417**,** 1-13.

NELSON, D. L., LEHNINGER, A. L. & COX, M. M. 2008. *Lehninger Principles of Biochemistry,* New York, W. H. Freeman.

NICHOLLS, D. G. & BUDD, S. L. 2000. Mitochondria and neuronal survival. *Physiol Rev,* 80**,** 315-60.

STARKOV, A. A., POLSTER, B. M. & FISKUM, G. 2002. Regulation of hydrogen peroxide production by brain mitochondria by calcium and Bax. *J Neurochem,* 83**,** 220-8.

THEUREY, P., CONNOLLY, N. M., FORTUNATI, I., BASSO, E., LAUWEN, S., FERRANTE, C., MOREIRA PINHO, C., JOSELIN, A., GIORAN, A. & BANO, D. 2019. Systems biology identifies preserved integrity but impaired metabolism of mitochondria due to a glycolytic defect in Alzheimer's disease neurons. *Aging cell,* 18**,** e12924.

WARD, M. W., HUBER, H. J., WEISOVA, P., DUSSMANN, H., NICHOLLS, D. G. & PREHN, J. H. 2007. Mitochondrial and plasma membrane potential of cultured cerebellar neurons during glutamate-induced necrosis, apoptosis, and tolerance. *J Neurosci,* 27**,** 8238-49.

WARD, M. W., REGO, A. C., FRENGUELLI, B. G. & NICHOLLS, D. G. 2000. Mitochondrial membrane potential and glutamate excitotoxicity in cultured cerebellar granule cells. *J Neurosci,* 20**,** 7208-19.

WEI, A. C., AON, M. A., O'ROURKE, B., WINSLOW, R. L. & CORTASSA, S. 2011. Mitochondrial energetics, pH regulation, and ion dynamics: a computational-experimental approach. *Biophys J,* 100**,** 2894-903.

YOSHIDA, T., KAKIZUKA, A. & IMAMURA, H. 2016. BTeam, a Novel BRET-based Biosensor for the Accurate Quantification of ATP Concentration within Living Cells. *Sci Rep,* 6**,** 39618.
